# A Brain Circuit for Status Epilepticus

**DOI:** 10.64898/2026.07.17.738338

**Authors:** Neal Nolan, Payam Tabaee Damavandi, Joseph I. Turner, Artemis Iatrou, Ross C. MacFadyen, William Drew, Arun Garimella, Bassam Al-Fatly, Clemens Neudorfer, Aaron E.L. Warren, Samuel B Snider, Andrew R. Pines, Natalia Rost, Ona Wu, Pilar Bosque-Varela, Lukas Machegger, Johannes A.R. Pfaff, Marian Galovic, Sattar Khoshkhoo, Urs Fisch, Jurriaan M Peeters, John D. Rolston, Andrew J. Cole, Jong Woo Lee, Eugen Trinka, Michael D. Fox, Giorgi Kuchukhidze, Frederic L.W.V.J. Schaper

**Author notes:** **Corresponding Authors:** Neal Nolan MD, 60 Fenwood Road, Boston, MA 02115, U.S.**, email:** and, Frederic L.W.V.J. Schaper MD PhD, 60 Fenwood Road, Boston, MA 02115, U.S.**, email:**. These authors contributed equally (Neal Nolan, Payam Tabaee Damavandi, Joseph I. Turner, Giorgi Kuchukhidze, Frederic L.W.V.J. Schaper).

## Abstract

Status epilepticus (SE) is a life-threatening persistent epileptic seizure that can arise from various brain structures, leaving its brain circuit unknown. In this study, we utilize brain imaging changes during SE to reveal the brain architecture and circuit of persistent seizures. Multimodal lesion mapping identified that brain imaging changes during SE localize to a specific predisposed brain architecture characterized by increased metabolic rate, high synaptic and mitochondrial density, glutamate (mGLUR5 and NMDA) and GABA receptors. Gene expression patterns within lesion locations revealed a transcriptomic profile enriched for epilepsy pathologies (including SE), neuronal and synaptic processes, and glutamate signaling. Lesion network mapping demonstrated these same lesions map to a common brain circuit, unifying a traditionally heterogeneous patient population. Findings were validated in an independent cohort and the identified SE circuit distinguished brain imaging changes during SE from other lesion etiologies with excellent accuracy (91%), significantly outperforming all other tested maps. With this SE circuit, we identify therapeutic targets for precision therapy that could modulate this circuit. This study demonstrates brain imaging changes in SE converge on a unified brain circuit that could help diagnostic workup of patients in critical care and guide clinical trials of precision therapy for persistent seizures.

## Introduction

Status epilepticus (SE) is a state of persistent epileptic seizure and a common neurologic emergency with poor response to antiseizure medications in up to 50% of cases.(*1–3*) SE can lead to irreversible brain damage,(*4, 5*) and refractory cases have a fatality rate approaching 40%,(*2, 6*) placing it among the most lethal of neurological emergencies. While epilepsy is recognized as a brain network disease and precision therapies increasingly target the seizure network,(*7, 8*) the brain network underpinning SE remains largely unknown.(*9–11*)

Mapping the brain network of SE poses several challenges as it is a transient brain state that can arise from various brain regions. To map a patient’s seizure network directly, epilepsy surgery teams often use intracranial EEG in the pre-operative setting. However, this is challenging to apply in the critical care setting of SE, including due to lack of a clear anatomo-electroclinical hypothesis to guide implantations.(*12–14*) In contrast, diffusion magnetic resonance imaging (MRI) is often used in SE clinical care and does not rely on an *a priori* neuroanatomical hypothesis.

Diffusion MRI can identify transient brain lesions during SE, also known as peri-ictal MRI abnormalities (PMA). PMA are associated with drug resistance, poor clinical outcome, and are temporally coupled to SE.(*15–18*). PMA are hypothesized to result from the intense metabolic demands of sustained seizure activity,(*19–21*) leading to cytotoxic edema visible as diffusion restricted brain lesions.(*15, 16, 19, 21, 22*) Due to their focal localization and temporal coupling, PMA could provide a ‘*snapshot’* of the brain network involved.(*23–25*) However, as previous studies localized PMA to disparate brain regions, their network localization remains unclear.(*15, 26, 27*) Mapping the brain structures and networks PMA intersect could thus help establish a neuroanatomical hypothesis of SE, distinguish seizure-related lesions from other lesion etiologies, and identify precision targets.

## Results

### Peri-ictal MRI abnormalities localize to brain areas high in brain metabolism and synaptic, mitochondrial, and neurotransmitter receptor density

To test whether PMA locations map to specific brain architecture (**Fig. 1, first row**), we used a systematic search of published case reports (**Supplementary Fig. 1**) and generated a discovery cohort (n=582, 92 [16%] with SE) of 92 SE case lesion locations (**Supplementary Fig. 2A**) and 490 stroke control lesion locations (**Supplementary Fig. 2B**). Demographics and details on the search strategy and selection of cases and control lesion locations can be found in **Supplementary Fig. 1** and **Supplementary Table 1**. We segmented the diffusion restricted lesion locations in template space (MNI 152) and tested for overlap of each lesion location with the Harvard-Oxford (sub)cortical atlas of brain regions,(*28*) an atlas of vascular territories,(*29*) and normative maps of brain microstructure, metabolism, and neurotransmitter systems. In total we tested 52 maps, of which 46 were derived from “NeuroMaps” from https://github.com/netneurolab/neuromaps and 6 from other sources such as HumanMitoBrainMap from http://humanmitobrainmap.bcblab.com and the Human Connectome Project from https://www.humanconnectome.org/study/hcp-young-adult/article/s1200-group-average-data-release)(30–32) (**Fig. 2A, D, G**). A detailed description of each map and its source can be found in **Supplementary Table 2**. In a data-driven manner, we identified 10 candidate maps that best distinguished SE vs. control lesion locations in the discovery cohort (highest *t-value)*, after Bonferroni correction for multiple comparisons across 52 maps (**Fig. 2B, E, H; Supplementary Fig. 3**). We then tested the top 10 maps for their predictive capacity to distinguish between SE vs. control lesions in the validation cohort (n=166, 46 [28%] with SE, **Fig. 2C, F, I**).

**Figure 1.**
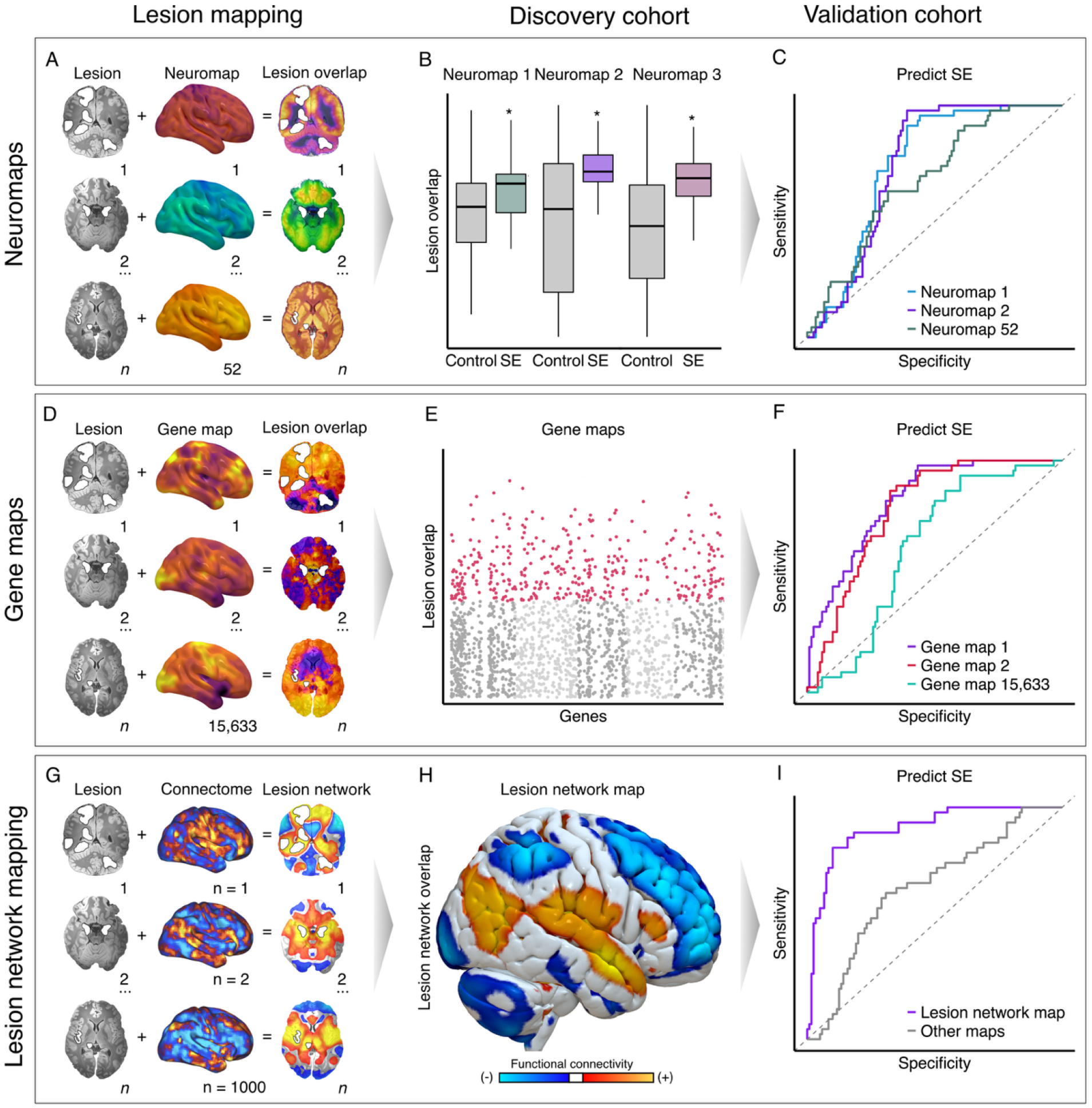
Study overview. Lesion locations of peri-ictal MRI abnormalities (PMA) and control lesions are overlapped on multiple atlas maps of brain microstructure, metabolism, and neurotransmitter receptors (left panels, second column) derived from Neuromaps, and MitoMap **(A)**. Lesion overlap is calculated and compared across SE and control groups by overlapping the lesion locations from the discovery cohort (n=582) with each atlas map **(B)**. Each data-driven map that distinguishes between SE and control groups in the discovery cohort is next tested for its capacity to predict SE versus control subjects in an independent validation cohort (**C**). The same analysis is performed for 15,633 maps of gene distributions from the Allen Human Brain Atlas (**D**). Candidate gene expression maps were identified in a data-driven manner in the discovery cohort (**E, each dot is a gene map**) and their predictive capacity was similarly tested in the validation cohort (**F**) Finally, the brain network connected to each diffusion restricted brain lesion in the discovery cohort is computed using lesion network mapping (**G**) and a lesion network map (**H**) was identified in a data-driven manner by overlapping each lesion network and assessing specificity versus control lesions. The lesion network map was likewise tested for its predictive capacity to distinguish between SE vs. controls in the validation cohort (**I**). *Abbreviations: SE, status epilepticus; DWI, diffusion-weighted imaging; PMA, peri-ictal MRI abnormality*.

**Figure 2.**
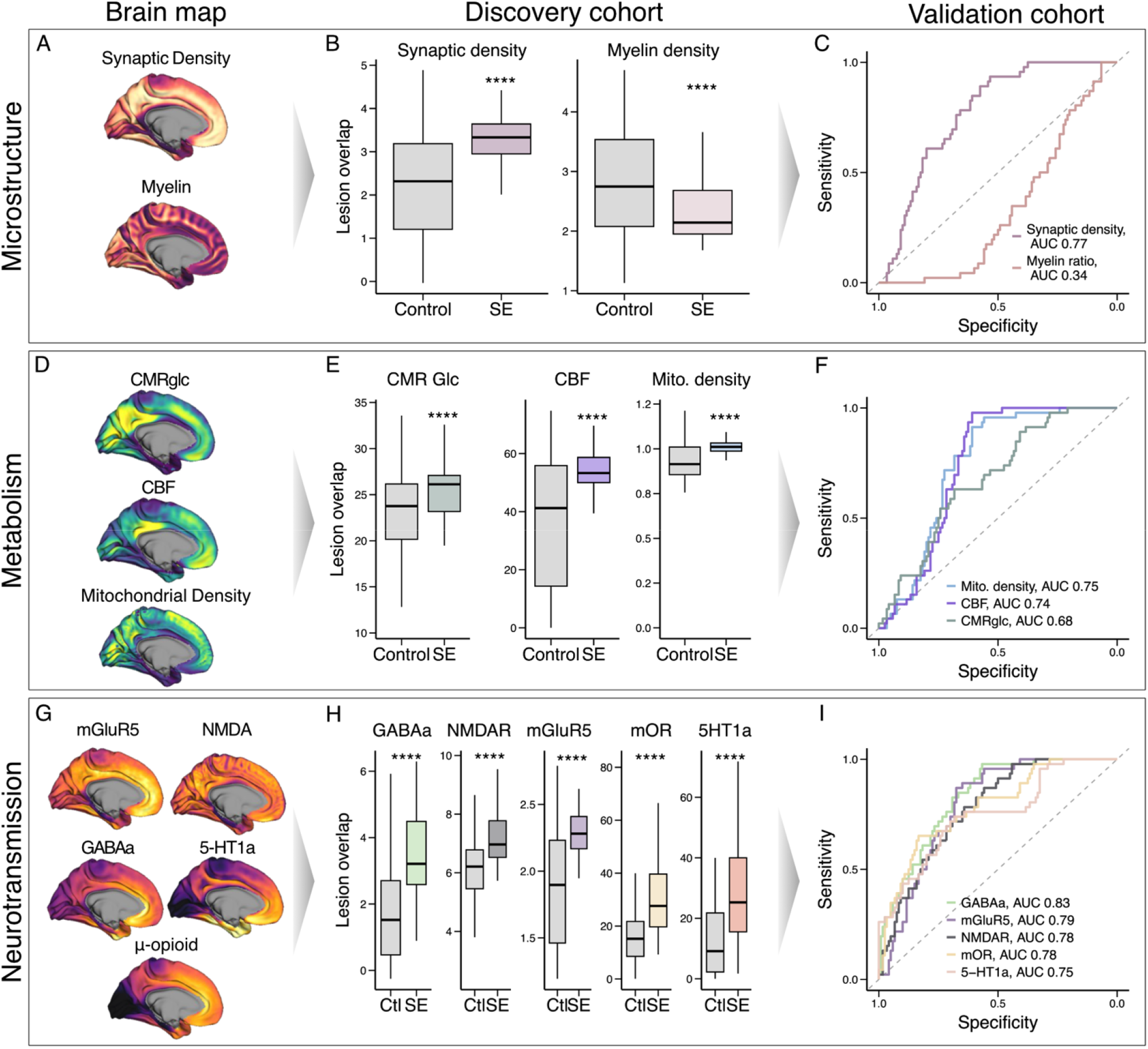
Lesion overlap with brain microstructure, metabolism, and neurotransmitter systems. Lesion locations of the discovery cohort were segmented in MNI152 space and overlapped on top of a collection of 52 atlas maps (**first column**). In a data-driven manner, we identified 10 candidate maps that distinguished between SE and control groups in the discovery cohort (**second column, Supplementary Table 2**). To assess each map’s capacity to predict SE we calculated lesion overlap of the validation cohort with each of the 10 candidate maps (**third column**). This process was repeated for maps of brain microstructure (**first row**), metabolism (**second row**), and neurotransmitter receptors (**third row**). Candidate maps of microstructure (synaptic density and myelin ratio, **A**) that distinguished between SE and control lesions in the discovery cohort (**B**) also predicted SE in the validation cohort (**C**). Candidate maps of metabolism (mitochondrial density, cerebral blood flow (CBF), cerebral metabolic rate of glucose (CMRgluc), **D**) that distinguished between SE and control lesions in the discovery cohort (**E**) also predicted SE in the validation cohort (**F**). Candidate maps of neurotransmitter systems (**G**) that distinguished between SE and control lesions in the discovery cohort (**H**) also predicted SE in the validation cohort (**I**). Lesion overlap on candidate maps was compared between SE and control groups with a two-sample *t*-test. Predictive capacity of each map was calculated with ROC analysis. **** denoting *p* < 0.0001 after familywise error correction for multiple comparisons across 52 maps. *Abbreviations: SE, status epilepticus; PMA, peri-ictal MRI abnormalities; AUC, area under the curve; CBF, cerebral blood flow; CMR Glcglucose cerebral metabolic rate*.

PMA occurred more frequently in the occipital (χ^2^ = 48.5, *p* < 0.001), temporal (χ^2^ = 36.8, *p* < 0.001), and parietal lobes (χ^2^ = 13.8, *p* = 0.008) compared to control lesions and were located across multiple vascular territories. PMA lesions more frequently intersected the posterior cerebral artery (PCA) territory compared to control lesions (χ^2^= 45.5, *p* < 0.001), while there was no difference in other vascular territories (**Supplementary Table 3**). We found PMA localized to brain regions high in synaptic density (*p* < 0.001) and low in myelin (*p* < 0.001) (**Fig. 2B**). Both maps of synaptic density (AUC = 0.77) and myelin (AUC = 0.66) significantly predicted SE versus control lesions in the validation cohort (**Fig. 2C**). Since previous anecdotal reports in patients with SE hypothesized that PMA result from the increased metabolic demands of sustained seizure activity,(*16–19, 21, 23, 26*) we investigated the spatial relationship between PMA locations and maps of brain metabolism. Indeed, PMA localized to regions high in cerebral blood flow (CBF, *p* < 0.001), glucose utilization (glucose cerebral metabolic rate, CMRglucose, *p* < 0.001), and mitochondrial density (*p* < 0.001, **Fig. 2E**). Highest predictive capability among these was found for CBF (AUC = 0.744), followed by mitochondrial density (AUC = 0.745) and then glucose utilization (AUC = 0.679, **Fig. 2F**). Finally, to investigate the relationship of PMA to specific neurotransmitter systems, we evaluated lesion intersection with maps of neurotransmitter receptors derived from PET tracer binding. We found PMA localize to regions with mGluR5 metabotropic glutamate (*p* < 0.001); NMDA (*p* < 0.001); GABA_A_ (*p* < 0.001); 5-HT1A receptors (*p* < 0.001); and μ-opioid receptors (MOR, *p* < 0.001, **Fig. 2H**). Highest predictive capability in the validation cohort was found for GABA_A_ (AUC = 0.828), followed by the mGluR5 (AUC = 0.791), μ-opioid (AUC = 0.784), NMDA (AUC = 0.777), and 5-HT1A (AUC = 0.747) receptors (Fig. 2I). In contrast, maps of dopamine and histamine receptor binding, 5-HT4 and 5-HT6 receptors, norepinephrine transporters and COX among others did not distinguish between SE and control patients (**Supplementary Fig. 3**).

### Peri-ictal MRI abnormalities localize to gene expression patterns enriched in epilepsy pathologies, glutamate signaling and neuronal and synaptic processes

To test whether PMA locations (**Supplementary Fig. 2**) map to a specific gene expression patterns (**Fig. 1, second row**), we calculated intersection of SE and control lesion locations of the discovery cohort (n=582, **Supplementary Fig. 2**) with normative maps of gene expression. We tested lesion overlap with a total of 15,633 gene distributions derived from the Allen Human Brain Atlas (AHBA, https://human.brain-map.org(33), **Fig. 1D-F**) Gene expression maps were generated using *abagen* (https://github.com/rmarkello/abagen) with recommended settings.(*33, 34*) In a data-driven manner, we identified the top 5 candidate gene maps associated with PMA in the discovery cohort (highest *t-*value), after Bonferroni correction for multiple comparisons across 15,633 maps. We subsequently tested each candidate map’s predictive ability to distinguish between SE and control lesions in the validation cohort. Among 15,633 gene expression patterns, the 5 that showed the strongest association with PMA locations (Bonferroni corrected *p value* < 0.05) were *HOMER2*, *CARTPT*, *GRM7*, *C1orf53*, and *CPNE4* (**Fig. 3A**). These same gene expression maps predicted SE lesion etiology in the independent validation cohort with considerable accuracy (AUC = 0.794 - 0.844, **Fig. 3B, C**). To identify the biological pathways associated with these gene expression patterns, the lesions in the discovery and validation cohorts were pooled and calculated intersection with each gene map (n = 15,633). Genes were then ranked based on the effect size (highest *t-value*) from the comparison of SE vs control lesions. Enrichment of terms from the Human Phenotype Ontology and Gene Ontology databases was evaluated using gene set enrichment analysis (GSEA).(*35*) PMA locations showed gene enrichment for epilepsy-related terms (*epilep\** or *seiz\**, P = 9.9e-05, **Fig. 3D**), multiple epilepsy-related pathologies including SE (**Fig. 3E**), and pathways related to neuronal and synaptic processes including neurotransmitter activity, synapse assembly, vesicle-mediated transport, and glutamate signaling (**Fig. 3F**). In contrast, stroke control lesion locations showed enrichment in non-neuronal processes including development and remodeling of the vasculature, immune and extracellular matrix components, vascular and inflammatory pathologies, as well as pathways related to cytotoxicity and ribosomal proteins (**Supplementary Fig. 4**).

**Figure 3.**
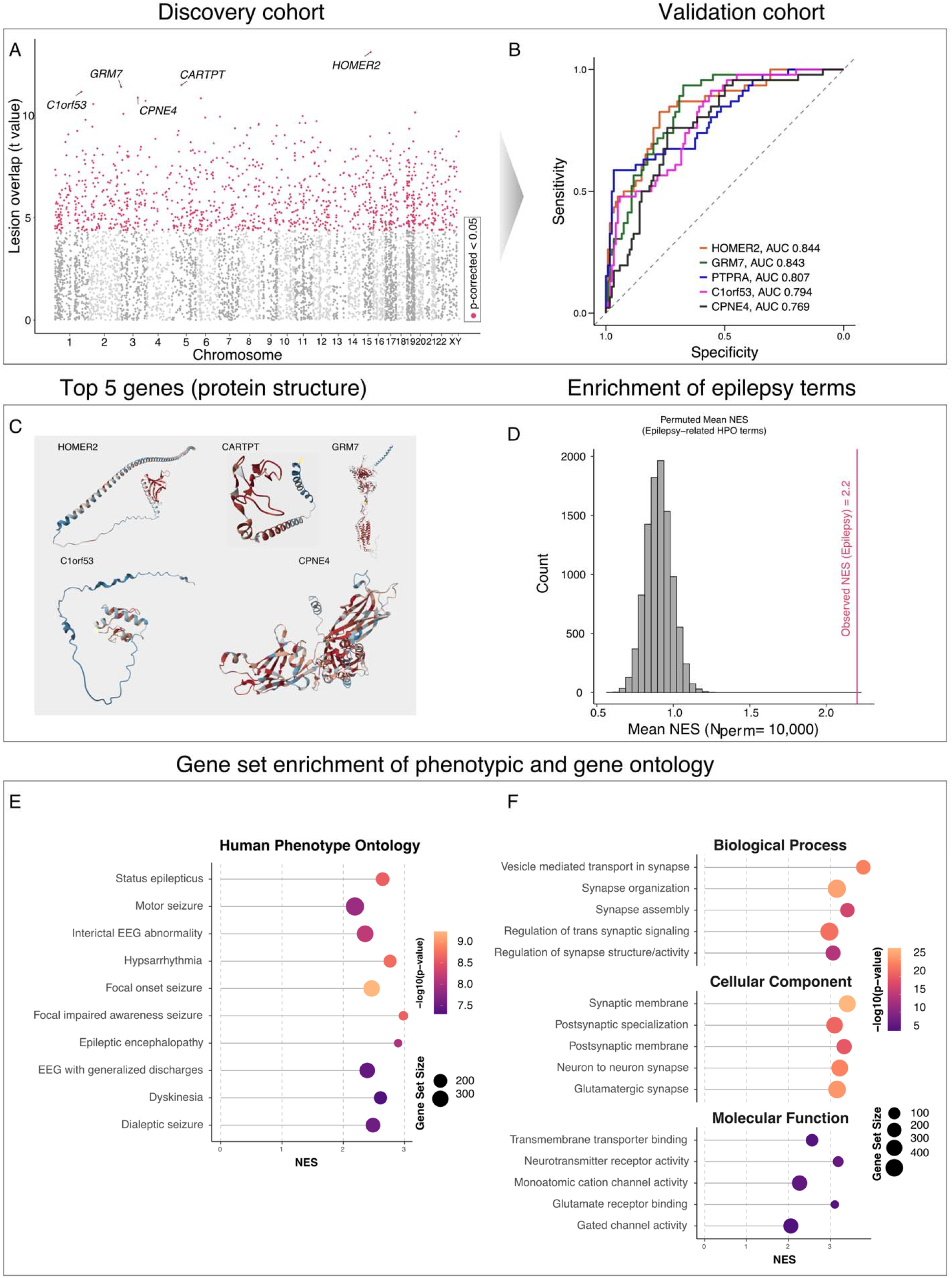
Lesion overlap with gene expression. Maps of gene expression were obtained from the Allen Human Brain Atlas (AHBA) and preprocessed with *abagen*. Lesion overlap of lesions from the discovery cohort with the gene expression maps of the 15,633 genes was calculated and compared across SE and control groups. In a data-driven manner, we identified the top 5 gene distributions best distinguishing between SE and controls groups. Panel **A** shows a Manhattan plot of the genes where SE lesion overlap exceeded control lesion overlap, with the pink colored dots denoting the genes with a Bonferroni corrected p-value < 0.05. The top 5 genes as determined by *t-value* are labeled. These top 5 gene distributions were then tested for their predictive capacity in distinguishing SE vs. controls in the validation cohort using ROC analysis (**B**). The protein structures of the top 5 genes (*Homer2, CARTPT, GRM7, C1orf53, CPNE4)* were derived from the *alphafold* database and illustrated (**C**). We then pooled the discovery and validation cohort and repeated the lesion overlap analysis, ranking each gene in their capacity to distinguish between SE and control groups (highest *t-value*). Gene set enrichment analysis (GSEA) was performed on this rank list using the Human Phenotype Ontology (HPO) database. Epilepsy-related terms showed higher enrichment than the null distribution of mean normalized enrichment scores (NES) across 10,000 permutations (**D**; red line). The top 10 most significantly enriched HPO terms **(E)** linked PMA lesions to epilepsy pathologies. Expansion of this analysis to the Gene Ontology Database (**F**) revealed the top 5 biological processes, cellular components and molecular functions including neuronal and synaptic processes. *Abbreviations: NES, Normalized enrichement score; HPO, Human Phenotype Detabase; AUC, area under the curve*.

### Lesion network mapping of peri-ictal MRI abnormalities identifies a unifying SE network

To test whether PMA locations (**Supplementary Fig. 2**) map to a common brain network (**Fig. 1, third row**), we performed lesion network mapping (**Fig. 1 G-I**).(*36–38*) In a data-driven manner, PMA and control lesions from the discovery cohort were run as seeds through a large normative functional connectome (n=1,000) to identify the brain network functionally connected to each lesion location. Lesion network overlap (or sensitivity) analysis revealed >85% of PMA locations were functionally connected to peak regions in the pulvinar nucleus of the thalamus and the precuneus cortex (discovery cohort, **Fig. 4A-B)**. Lesion network specificity analysis revealed PMA locations were more functionally connected to these same regions compared to control lesions (family-wise error corrected P < 0.05, **Fig. 4C**). For specificity analyses, we used 10,000 label permutations to identify functional connections specific to SE vs. control lesion locations and corrected for multiple comparisons across all brain voxels using fwe correction. Label permutation ensures that identified connections are specific to epilepsy and not driven by non-specific properties of the human connectome.(*39–41*). Key connections both sensitive and specific to SE were identified by retaining voxels that had the same sign across sensitivity and specificity analyses followed by multiplying the values of these shared voxels.(*42, 43*) The resulting ‘agreement map’ of peak sensitivity and specificity on a voxel-wise basis is hereafter termed an ‘SE network’ (**Fig. 4C, right; Fig. 4D**) and was tested for its predictive capability in distinguishing SE vs. control lesions in the validation cohort.

**Figure 4.**
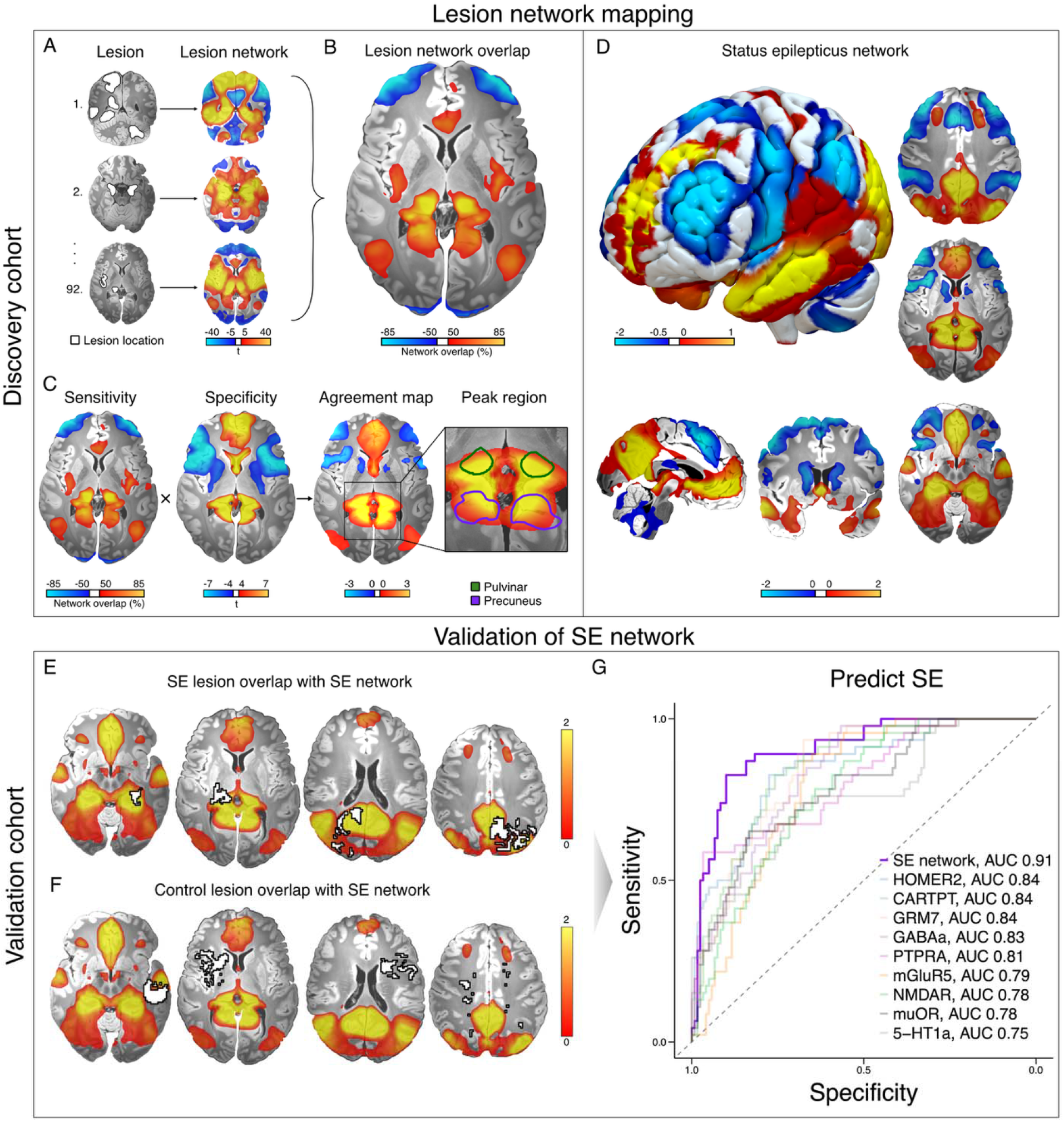
Lesion network mapping of peri-ictal MRI abnormalities and identification of a SE network. PMA locations (**A, left**) and the brain network functionally connected to each PMA location was identified using the human connectome, resulting in a lesion network for each individual subject **(A, right)**. Lesion networks were overlapped, identifying common functional network connections across PMA in the pulvinar nucleus of the thalamus and precuneus cortex (peak sensitivity 85%, **B**). These same regions (**C, left**) were also specific for SE compared to lesion networks of control subjects (*p* < 0.05, **C, middle**). Agreement mapping identified an SE network comprising voxels sensitive and specific to SE with peaks in the medial pulvinar nucleus of the thalamus and precuneus cortex (**C, right; D**) with strong connectivity to the posterior cortex, ventromedial prefrontal cortex, cingulate, hippocampus, claustrum, septum, midbrain, tectum, and cerebellum. Lesion overlap of lesion locations from the validation cohort with the SE network derived from the discovery cohort demonstrates that PMA fall predominantly within the SE network (**E**) while control lesion locations predominantly fall outside the SE network (**F**). The SE network was able to distinguish PMA from control lesion locations in an independent validation cohort (**G**) with excellent accuracy (AUC_SE network_ = 0.906, *p* < 0.001), significantly outperforming the next best predictive map *HOMER2* (DeLong test, paired, Z = 2.44, *p* = 0.015). *Abbreviations: SE, status epilepticus; PMA, peri-ictal MRI abnormalities; AUC, area under the curve*.

The SE network included positive functional connectivity to the pulvinar nucleus of the thalamus, precuneus cortex, mesial frontal lobe, lateral temporal poles, posterior parietal lobe, splenium of the corpus callosum, posterior insula, claustrum, septal nuclei, midbrain, tectum; and negative connectivity to the precentral gyrus, dorsolateral prefrontal cortex, anterior parietal lobe, anterior insula, cerebellum, and medulla. (**Fig. 4D**) We then tested the SE network’s predictive capacity to distinguish SE from control lesions in the validation cohort. In the validation cohort, lesion intersection with the SE network derived from the discovery cohort predicted SE vs. control (**Fig. 4E,F**) with excellent accuracy (AUC_SE network_ = 0.91, **Fig. 4G**). The SE network reached a sensitivity of 83% and specificity of 90% at Youden’s *J* balanced cutpoint (**Supplementary Fig. 5**), significantly outperforming the next strongest predictive map (*HOMER2* gene expression) and all other tested maps (Delong test, paired, Z = 2.44, *p* = 0.015). The SE network was robust across different control and subgroup analyses and independent of properties of the human connectome (**Supplementary Results A**).(*39–41*)

### The SE network identifies testable precision targets

Because prior work found lesion locations can be used to map therapeutic targets for brain stimulation in movement(*42, 44–47*) and mood disorders,(*48–50*) the SE network may have the potential to identify ‘*chokepoints*’ or therapeutic targets to interrupt persistent seizures. To identify these targets, we analyzed a previously published precomputed connectome(*48, 51*) and identified the voxels whose whole-brain functional connectivity profile best resembled the SE network. In the thalamus, we found a collection of voxels in the (ventral medial) pulvinar nucleus (MNI -12 -35 1) whose connectivity profile spatially correlated with the SE network (R = 0.66). To illustrate how this SE network peak may guide therapeutic interventions (**Fig. 5A**), we used Lead-DBS software (https://www.lead-dbs.org) to plan a mock lead trajectory targeting this location.(*52*) The same analysis was performed on voxels in the frontal cortex – a safe and effective transcranial magnetic stimulation (TMS) target used to treat mood disorders. We identified a collection of voxels in the medial prefrontal cortex (BA10) that spatially correlated with the SE network (R = 0.47) and used SimNIBS software (https://simnibs.github.io/simnibs/build/html/index.html) to plan the theoretical optimal positioning of a figure-of–eight TMS coil to target this location (MNI -2, 64, -4), approximately 1 cm below the midpoint between AFz and Fpz (**Fig. 5B**). Finally, we used the Allen Human Brain Atlas to identify a gene target whose expression pattern best resembled the SE network. We found that the top 1 out of 15,633 gene expression patterns best resembling the SE network (R = 0.30) was *WNT3* (**Fig. 5C**).

**Figure 5.**
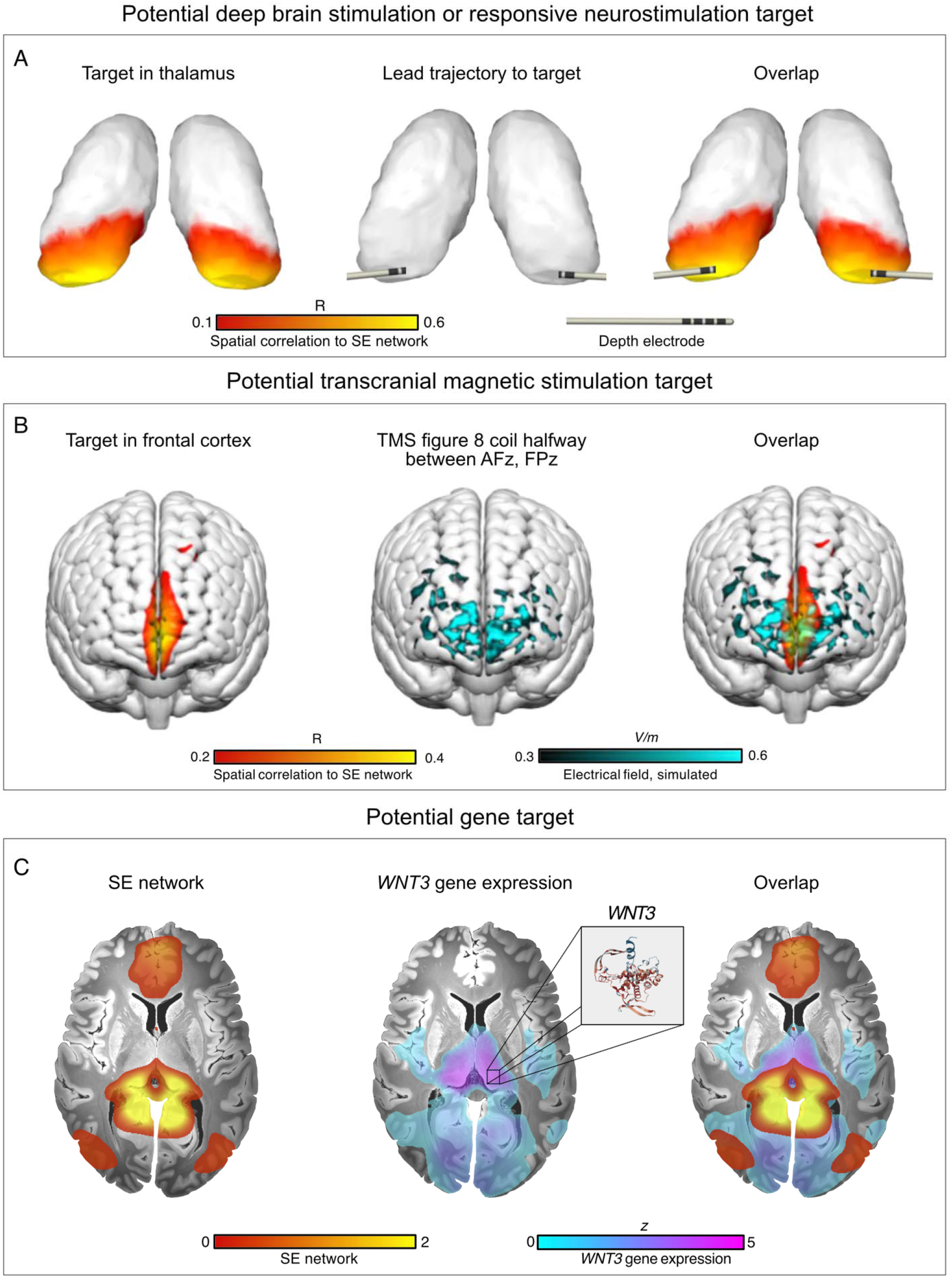
The SE network identifies testable therapeutic targets. With a precomputed connectome, we identified a peak in the ventral medial pulvinar nucleus whose whole brain connectivity profile was most representative of the SE network. For illustrative purposes and to aid potential clinical translation, we planned a mock depth lead trajectory to this potential thalamic target (**A**). With the same precomputed connectome, we identified a peak in the medial prefrontal cortex that best aligned with a TMS coil positioning between AFz and FPz (**B**). Finally, using the Allen Human Brain Atlas, we identified the top 1 gene out of 15,633 genes that best aligned with the SE network: *WNT3* (**C**). *Abbreviations: SE, status epilepticus; DBS, deep brain stimulation; RNS, responsive neurostimulation; TMS, transcranial magnetic stimulation*.

## Discussion

This study demonstrates brain imaging changes during SE localize to a specific predisposed brain architecture and map to a common brain network, unifying a traditionally heterogeneous patient population. These findings establish a neuroanatomical hypothesis of SE which could aid differential workup of patients in critical care settings and which identifies precision targets.

SE is a common stroke mimic(*53*) that can induce diffusion restriction on MRI.(*27*) Distinguishing diffusion restriction related to SE from restriction due to other etiologies can be time-sensitive in critical care scenarios involving unexplained brain lesions and an inconclusive EEG, for example in patients with non-convulsive SE.(*54*) Mapping the diffusion restricted brain locations specific to sustained seizure activity could aid differential workup in these settings. Consistent with prior studies localizing PMA, (*23, 24, 55, 56*) we find heterogeneous involvement of multiple regions in the cortex, splenium, hippocampus, thalamus, and cerebellum across 92 cases of SE. Why PMA frequently involve these distributed brain regions remains largely unknown.

Despite a heterogeneous involvement of distributed brain regions, we found PMA consistently localize to brain structures with a high synaptic but low myelin density. These findings are consistent with literature showing highly myelinated brain areas have an excitatory-inhibitory ratio shifted towards inhibition which functions as a “brake” on neuroplasticity,(*57*) a key proposed mechanism of seizure generation and epilepsy.(*58*) Likewise, areas low in myelin may be more predisposed to seizure, since a loss of this “brake” could shift the brain from inhibition towards excitation.(*57*) Consistent with these results, we find PMA localize to areas high in glutamate (mGluR, NMDA), GABA, serotonin and opioid receptors, which have all been implicated in the modulation excitatory-inhibitory balance, cortical excitability, and seizures.(*59–77*) Next, we tested the *a priori* hypothesis derived from previous clinical observations that regions of increased metabolic rate are more vulnerable to sustained seizures.(*19, 78, 79*) We found robust support for this hypothesis in our data, suggesting brain areas with a high metabolic rate at rest are predisposed to the intense metabolic demands of sustained seizure activity. In future therapeutic studies, these areas may be monitored during or following SE for signs of hypermetabolic neuronal injury,(*19, 21, 23*) irreversible cell loss, gliosis, or atrophy(*80–82*) which could inform development of biomarkers to monitor seizure-induced brain damage. On a gene expression level, we found that PMA follow gene expression patterns enriched in epilepsy-related neuronal processes such as synapse organization, vesicle-mediated transport, cation channel activity, and glutamate signaling, in particular in the metabotropic signaling receptor *GRM7* (mGluR7) and the key glutamatergic postsynaptic scaffolding protein *HOMER2* (Homer2).(*83, 84*) These findings suggest that PMA are not merely a result of the intense metabolic demands of sustained seizure activity but are enriched in brain areas with a gene expression pattern intrinsic to epilepsy pathology.

Using a method termed lesion network mapping,(*36, 37, 85*) we found evidence for a common brain network that connects disparate diffusion restricted brain lesions during SE across a heterogeneous population of patients, reconciling the varied results reported in prior studies.(*23, 26, 27*) This network includes key nodes in the pulvinar thalamus, claustrum, septum, cerebellum and regions of the limbic system and default mode network. Intersection of independent PMA locations with this ‘SE network map’ (**Fig. 4**) was able to distinguish diffusion restricted lesions related to SE from other lesion etiologies with excellent accuracy (AUC = 0.91, sensitivity of 83% and specificity of 90%). This SE network was an independent and significantly stronger predictor of SE than any single cortical lobe, vascular territory, brain region, metabolism or gene expression map. These unifying results were independent of seizure onset location, suggesting that despite the heterogeneity in seizure onset, SE ‘*hijacks*’ a preserved brain network, unifying a traditionally heterogenous patient population. A unifying brain network for SE establishes a testable hypothesis on the anatomo-electroclinical correlation of SE and may guide precision therapies to interrupt persistent seizures. Our results specifically point towards the pulvinar nucleus of the thalamus being an important node or ‘*chokepoint*’ of persistent seizures. The pulvinar is hypothesized to function as a subcortical relay in SE(*86*) and is a known node for seizure spread in both temporal and extratemporal seizures.(*87, 88*) Our results support this hypothesis and extend it by showing that even without apparent diffusion restriction, the pulvinar may still serve a central role in SE due to its intrinsic brain connectivity pattern. Several case series have reported the use of DBS of either the centromedian or anterior thalamic nuclei in patients with SE with mixed results.(*89, 90*) Our findings suggest the ventromedial pulvinar deserves additional attention, and we here propose a testable target for future studies. Although more speculative, we additionally identify a potential cortical network target for future transcranial magnetic stimulation (TMS) studies and potential gene targets such as *WNT3*, modulation of which has been shown in preclinical models to reduce neuronal excitability, regulate synaptic plasticity(*91–94*) and modulate seizures in models of induced SE.(*95–99*)

Several limitations are worth highlighting. First, lesions in the discovery cohort were segmented from 2D images available in the published literature consistent with previous lesion network mapping studies, which inherently do not reflect the total 3D lesion volume which was available for the independent validation cohort. Second, reporting of clinical details of published cases was variable, which adds noise to the data biasing against unifying findings. Third, it is important to highlight that the multimodal brain atlases used in this study were derived from normative data and not patients with SE. This was intentional as the aim of this study was to map the brain structures and networks PMA intersect and predispose to sustained seizure activity in the average human brain; not to identify how the brain changes during or following SE. Fourth, gene expression levels across brain regions do not always correspond to those genes’ functional importance. Finally, due to our retrospective study design, any clinical inferences should be interpreted with caution pending further controlled clinical studies.

This study demonstrates brain imaging changes during SE converge on a common brain network, unifying a heterogeneous patient population. Intersection of lesion locations with this SE network distinguishes brain imaging changes during SE from those due to other etiologies, which could help diagnostic workup of patients in critical care settings. This SE circuit could guide clinical trials of precision therapy to stop persistent seizures.

## Methods

This study was carried out in accordance with the Declaration of Helsinki, approved by the institutional review board of the Brigham and Women’s Hospital, Boston, Massachusetts, and exempted from obtaining informed consent based on the secondary use of research data (Protocol number 2020P002987). For the discovery cohort, preferred Reporting Items for Systematic Reviews and Meta-Analyses (PRISMA) guidelines were followed to identify published cases of SE-related PMA. For the independent validation cohort, the Ethics Committee of the Region of Salzburg, Austria approved the study on human subjects (approval number 415-E/2422).

### Discovery cohort

Cases of brain lesions during SE (n = 92), also termed peri-ictal MRI abnormalities (PMA), were selected from Grillo *et al.,* a systematic review that collated published case reports of seizure-related MRI abnormalities.(*27*) We selected only cases with (a) a diffusion restricted brain lesion visible on a published image of the brain MRI and (b) a diagnosis of SE according to established International League Against Epilepsy (ILAE) criteria at the time of publication.(*100*) A detailed description of the search strategy and case selection can be found in **Supplementary Fig. 1**. Identical to previous lesion network mapping studies,(*36, 101*) lesion locations of PMA (i.e. the hyperintense signal on the MR diffusion sequence) were segmented as a 2D slice on the MNI-152 brain template (2 x 2 x 2 mm resolution) using 3D Slicer software (https://www.slicer.org/).(*36, 101, 102*) In cases of multifocal PMA, lesion masks were added, binarized, and further analyzed as a single lesion. Locations of diffusion restricted brain lesions in patients with ischemic stroke without seizures were used as control lesion locations (n = 490).(*103*) Patient characteristics such as age, sex, lesion etiology, seizure semiology, and focal seizure onset zone as determined by scalp EEG were collected and summarized in **Supplementary Table 1**. Lesion overlap with cortical lobes, subcortical regions, and vascular territories was assessed using atlas masks derived from the Harvard-Oxford (Sub)Cortical Atlas(*104, 105*) (masks were thresholded and binarized at 25% probability), and the ‘Vascular Territory template and atlas in MNI space’(*106*) identical to a previous study (**Supplementary Table 3**).(*11*)

### Independent validation cohort

An independent validation cohort(*23*) of patients with diffusion restricted brain lesions associated with SE or other etiologies was used to test the generalizability of our findings. This independent cohort consisted of a collection of prospectively collected diffusion restricted brain lesions, of which 46 were from patients with SE diagnosed according to current ILAE criteria(*100*), and 120 from patients with ischemic stroke, intracranial hemorrhage, tumors, or encephalitis without seizures. Patient characteristics and imaging are summarized in **Supplementary Table 5** as well as described in greater detail in Bosque Varela *et al.*.(*23*) Lesion locations of PMA were segmented as a 3D volume in native space using each patient’s MR diffusion sequence and 3D Slicer software (https://www.slicer.org/). The initial tracing was performed by one investigator (PTD) and subsequently validated by a second investigator (LM). Hyperintensities on diffusion imaging were delineated slice-by-slice on the axial view, creating a 3D volume for each lesion location in the patient’s native MRI space. A single diffusion sequence could contain multiple hyperintense sites, each corresponding to a distinct lesion. Segmentations of all diffusion restricted lesion locations in a single patient were added, binarized and combined into a single lesion mask. Lesion masks were subsequently normalized to MNI-152 space with Advanced Normalization Tools (ANTs, http://stnava.github.io/ANTs/).(107)

### Lesion overlap with brain structure and function, metabolism, and neurotransmitter systems

To test whether PMA locations map to specific brain architecture, we overlapped the lesion masks of the discovery cohort (n=582) with a priori maps of brain microstructure, metabolism and neurotransmitter systems. Lesion overlap was calculated on the original (unthresholded) volume maps by averaging the reference map’s voxel values within a certain lesion mask (sum of values / number of voxels). Lesion overlap was compared between SE and control groups using a two-sample *t-test.* P-values were calculated with 10,000 permutations of the group labels and a Bonferroni corrected two-sided *p* < 0.05 was considered significant. We tested lesion overlap of the discovery dataset with a collection of 46 available volume-space maps within “NeuroMaps” from https://github.com/netneurolab/neuromaps. Neuromaps is an open-source resource that includes normative maps of PET tracer binding to different neurotransmitter receptors and maps of synaptic density, myelin water fraction, glucose metabolism, and cerebral blood flow among others. We tested 6 additional reference maps including mitochondrial density within HumanMitoBrainMap from http://humanmitobrainmap.bcblab.com and T1w/T2w fraction computed with the MR images of the Human Connectome Project from https://www.humanconnectome.org/study/hcp-young-adult/article/s1200-group-average-data-release.(30–32) For a detailed description of each reference map and its source see **Supplementary Table 2**. We calculated lesion overlap of each lesion in the discovery cohort (n=582) with each reference map and identified the top reference maps (highest *t-value)* from each category (brain structure, metabolism, neurotransmitter systems, **Supplementary Fig. 2**) in a data-driven manner. We then tested the top candidate maps for their predictive capacity to distinguish between SE vs. control lesions in the validation cohort (n=166) using Receiver Operating Characteristic analysis (ROC) and calculated the area under the curve (AUC).(*23*) A two-sided *p* < 0.05 was considered significant.

### Lesion overlap with gene expression patterns

To test whether PMA locations map to specific gene expression patterns, we overlapped the lesion locations of the discovery cohort (n=582) with maps of gene expression. We tested lesion overlap with a total of 15,633 gene distribution maps derived from the Allen Human Brain Atlas (AHBA, https://human.brain-map.org)(33). The AHBA is a normative atlas of gene expression across the human brain derived from post-mortem microarray analysis of 6 healthy individuals sampled across 3,702 brain regions. (*33, 108*) Voxel-wise brain maps of gene expression were generated for all 15,633 genes in the AHBA using recommended preprocessing settings in abagen (https://github.com/rmarkello/abagen) including intensity-based filtering of 0.5, differential stability probe selection, nearest-neighbor interpolation with donor-wise averaging, cross-donor robust sigmoid normalization, and voxel-wise z-scoring.(*33, 34, 109*) Lesion overlap was calculated in an identical manner as described above. P-values were calculated with 10,000 permutations of the group labels and a Bonferroni corrected two-sided *p* < 0.05 was considered significant. In a data driven manner, we calculated lesion overlap of each lesion in the discovery cohort (n=582) with each gene expression map (n=15,633) and identified the top 5 candidate gene maps that best distinguished between SE vs. control lesion locations (highest *t-value)*. We subsequently tested the top 5 gene candidates for their predictive capacity to distinguish between SE and control lesions in the validation cohort (n=166). ROC analysis was performed and the AUC was calculated. A two-sided *p* < 0.05 was considered significant. To identify the biological pathways associated with gene expression within PMA locations we performed Gene Set Enrichment Analysis (GSEA).(*35*) We pooled the discovery and validation cohorts and compared lesion overlap of SE and control lesions on each individual gene map (n = 15,633, thresholded *z* > 0). All 15,633 genes were then ranked based on effect size in distinguishing between groups (highest *t-value*). GSEA with adjusted settings (gene set size = 20-500) was subsequently performed on the ranked *t*-values using the Human Phenotype Ontology (HPO) and Gene Ontology (GO) databases from GSEA.(*35*) The HPO database provided gene sets associated with diagnoses, while the GO provided gene sets associated with biological processes, cellular components and molecular functions. To define epilepsy-related terms, we selected all HPO terms containing the strings *‘epilep’* or *‘seizure’* in the pathway name and labeled them as “epilepsy-related” (N = 40); all remaining terms were labeled “non-epilepsy” (N = 2,569). We then calculated the observed mean normalized enrichment score (NES) across epilepsy-related terms and assessed whether this mean was higher than expected by chance. To evaluate significance, we used a permutation-based approach where the 40 terms labeled as epilepsy-related were randomly reassigned across all HPO terms 10,000 times. For each permutation, we recalculated the mean NES, generating a null distribution. We computed an empirical *p*-value as the proportion of permutation-based mean NEAS values that exceeded the observed mean NES.

### Lesion network mapping and identification of a SE network

To test whether PMA locations map to a common brain network, we performed lesion network mapping consistent with published methods(*36, 101, 102*) Lesion networks were generated for each lesion location in the discovery cohort (n=582) using seed-based functional connectivity analyses with the resting state functional connectivity data (2 × 2 × 2 mm resolution) of 1000 healthy participants from the Brain Genomics Superstruct Project (GSP1000 connectome).(*110*) Preprocessing of these scans has been fully described elsewhere(*111, 112*) and included regression of noise variables derived from motion, CSF, white matter, and the global signal. As in prior studies, Fisher’s *<u>r</u>*-to-*z* transformed Pearson correlation values were computed between each lesion location and all other brain voxels. A one-sample t-test was used to identify the voxels that were significantly connected to each lesion location (ie: showing correlated fMRI signal fluctuations) resulting in a lesion network *t*-map. Lesion network *t*-maps of SE-related PMA cases (n = 92, **Supplementary Table 1**) were then thresholded at a t > 5 in both the positive and negative direction, binarized, and overlapped across all SE subjects to identify common network connections sensitive to SE.(*102, 113*) To identify network connections specific to SE, we compared the lesion networks of SE-related PMA (n = 92) to the lesion networks of control ischemic stroke subjects (n=490, **Supplementary Table 1**) using a whole-brain voxel-wise *t*-test implemented in the software Permutation Analysis of Linear Models (PALM) (https://fsl.fmrib.ox.ac.uk/fsl/fslwiki/PALM).(114–116) Significance was assessed using 2000 permutations of the group labels at each brain voxel and family-wise error correction for multiple testing. We then identified the voxels both sensitive and specific to SE lesions by computing the product of these two maps after *z*-scoring, including only those voxels who shared the same sign across both maps (positive or negative correlation). The resulting “agreement map” of both sensitive and specific voxels is hereafter referred to as the “SE network”.(*102*) We used the SE network as an ‘*a priori* atlas’ of brain regions predisposing or vulnerable to PMA and hypothesized that SE lesions fell within this atlas while control lesions fall outside this atlas. Lesion overlap was computed as described above. We tested the SE network for its predictive capacity to distinguish between SE and control lesions in the validation cohort (n=166), computed the AUC, and compared it to the AUCs of the top 9 previously tested maps using a paired DeLong test.(*117*)

### Control and subgroup analyses

To test the robustness of the SE network, we performed several control and subgroup analyses. Since PMA overlapped with several cortical lobes and vascular territories (**Supplementary Table 3**), we used the validation cohort to evaluate whether lesion overlap with the SE network was an independent predictor of SE while controlling for each cortical lobe and the PCA vascular territory. Univariate and multivariate logistic regression models were computed predicting SE vs. controls including (i) lesion overlap with the SE network, (ii) cortical lobes or (iii) PCA vascular territory as independent variables. Odds ratios were calculated and each model’s performance was evaluated by computing the McFadden pseudo-R^2^. (**Supplementary Table 4**) Since PMA can both include cortical and subcortical brain lesions, we evaluated whether our results were driven by either cortical or subcortical involvement. To test this, we computed ROC curves in subgroups of patients with either (i) only cortical, (ii) cortical and subcortical, or (iii) only subcortical PMA. (**Supplementary Fig. 5**). As PMA frequently intersected the pulvinar or precuneus (53% across the discovery and validation dataset), we likewise tested whether predictive capacity was exclusive to pulvinar or precuneal lesions. We therefore excluded all patients (SE and control) with pulvinar or precuneal lesion involvement and recomputed the ROC curves of the validation cohort (**Supplementary Fig. 6**). To assess whether the SE network derived from our discovery cohort was exclusive to any specific focal seizure onset zone, we regenerated the SE network in subgroups of patients with either frontal, temporal or posterior cortex seizure onset (as defined by scalp EEG) and tested each of these networks separately for their predictive capacity in the validation cohort (**Supplementary Fig. 7**). We performed a similar analysis in subgroups of patients with convulsive or non-convulsive SE semiology (**Supplementary Fig. 8**). A recent study proposed that lesion networks may share spatial properties due repeated sampling of average brain connectivity in the human connectome.(*39*) We therefore regenerated the SE network controlling for various measures of average brain connectivity (referred to as Sum of C in van den Heuvel *et al.* 2026(*39*) and recomputed the ROC curves predicting SE in the validation cohort (**Supplementary Fig. 9, 10**).

### Identification of potential therapeutic targets

To identify potential therapeutic targets that may modulate the SE network, we used a precomputed connectome to identify the voxels most connected (i.e. whose whole brain connectivity profile is most similar) to the SE network. ^52,53^ In short, the connectivity profile of each brain voxel was computed by seed-based functional connectivity analysis (Pearson’s R at each brain voxel) using the preprocessed fMRI data of 1000 healthy participants from the Genomics Superstruct Project (GSP connectome).(*110*) Each voxel’s connectivity profile was then spatially compared to the SE network using a spatial correlation (Pearson R), creating a heatmap of voxels most connected to this network, referred to as a SE network target map. Peaks in this SE network target map were identified within the thalamus, as it is a safe and frequently used DBS and RNS target, and in the frontal cortex, as it is a safe and frequently used TMS target. First, using Lead-DBS software (https://www.lead-dbs.org), we illustrate a mock depth lead trajectory in the peak of the SE network target map within the ventral medial pulvinar nucleus (MNI 12, 30, 2). A lateral entry point in the superior temporal gyrus was chosen akin to a conventional SEEG lead trajectory. Second, using SimNIBS software (https://simnibs.github.io/simnibs/build/html/index.html), we illustrate a mock figure-of–eight TMS coil positioning at the peak of the SE network target map within the frontal cortex along the midline of the frontal poles (MNI -2, 64, -4). Third, using the Allen Human Brain atlas (https://human.brain-map.org), we illustrate a potential gene target by spatially correlating (Pearson’s R) each of the 15,633 gene expression maps to the SE network.

## Supporting information

Supplementary Results Figures and Tables

## Author Contributions

Conception and design of study: NN and FLWVJS

Design of analytical procedures: NN, JT, AI, and FLWVJS

Data contributors: NN, PTD, OW, NR, BAF, CN, AW, SBS, ARP, PBV, JARP, JP, MG, LM, JR, UF, JMP, JDR, AJC, JWL, ET, MDF, GK, JT, FLWVJS

Preprocessing and preparation of data for neuroimaging analysis: NN, PTD, WD, JT, RM, BAF, CN, PBV, GK, AG, SK and FLWVJS

Neuroimaging, genetic, and statistical analyses: NN, JT, AI and FLWVJS

Interpretation of data and analyses: NN and FLWVJS, with input from MDF, GK

Writing of manuscript, figures: NN and FLWVJS, with input from all authors

## Conflicts of Interest

NN reports personal fees as an employee of Rapport Therapeutics unrelated to the submitted work. JP reports personal fees from Boehringer Ingelheim AG & Co. KG, Medtronic, Eli Lilly, CERENOVUS (Johnson & Johnson Medical Products GmbH), research funding (directly, or to his institution) from Vesalio, all outside the submitted work. MG received fees from Advisis, Angelini Pharma, Bial, Eisai, Nestlé Health Science, and UCB outside the submitted work. SK is cofounder and advisor of Mosaica Medicines. JL has received grant support from NH/NINDS, SK Biopharmaceuticals, and UCB; he is a co-founder of Soterya, Inc. ET has received consultancy fees from Arvelle Therapeutics, Argenx, Clexio, Celegene, UCB Pharma, Eisai, Epilog, Bial, Medtronic, Everpharma, Biogen, Takeda, Liva-Nova, Newbridge, Sunovion, GW Pharmaceuticals, and Marinus; speaker fees from Arvelle Therapeutics, Bial, Biogen, Biopass, Böhringer Ingelheim, Eisai, Everpharma, GSK, GW Pharmaceuticals, Hikma, Liva-Nova, Newbridge, Novartis, Sanofi, Sandoz and UCB Pharma; research funding (directly, or to his institution) from GSK, Biogen, Eisai, Novartis, Red Bull, Bayer, and UCB Pharma outside the current study. ET is the Codirector of the European Consortium on Epilepsy Trials and Co-Founder of PrevEp Inc. MDF is a scientific consultant for Magnus Medical. MDF has intellectual property on the use of brain connectivity imaging to analyze lesions and guide brain stimulation, has consulted for Magnus Medical, Soterix, Abbott, Boston Scientific, Tal Medical, MDC Venture Capital, and is on the Scientific Advisory Board of Salma Health. He has received research support from Neuronetics and Boston Scientific. PBV received travel grants and honoraria from UCB, not related to the presented work.

## Funding

FLWVJS was supported by the National Institutes of Health (R01NS127892) and American Epilepsy Society (846534). SK was supported by the National Institutes of Health (K08NS128272) and Career Award for Medical Scientists from the Burroughs Wellcome Fund. G.K. was supported by FWF (Fonds zur Förderung der wissenschaftlichen Forschung), Austrian Science Fund; Project number KLI 969-B. E.T. was support by grants from the Austrian Science Fund (FWF), Österreichische Nationalbank, and the European Union. MDF was supported by grants from the NIH (R01MH113929, R21MH126271, R21NS123813, R01NS127892, R01MH130666, UM1NS132358), the Kaye Family Research Endowment, the Ellison / Baszucki Family Foundation, the Once Upon a Time Foundation, The Milken Foundation / BD2, the Manley Family, Donna and Tom May, and Chuck and Kerri Bean. JDR was supported by the National Institutes of Health (R01NS136297) and the Once Upon a Time Foundation.

