## Supplementary Results Figures and Tables for "A Brain Circuit for Status Epilepticus"

### Supplementary Materials

**Supplementary Results:**

Supplementary Results A: The SE network is robust across different control and subgroup analyses and is independent of properties of the human connectome

**Supplementary Figures:**

Supplementary Figure 1: PRISMA search strategy

Supplementary Figure 2: Status epilepticus and control brain lesions

Supplemental Figure 3: Lesion overlap of SE vs. controls in the discovery cohort on maps of brain structure, metabolism, and neurotransmitter receptors

Supplemental Figure 4: Gene expression patterns enriched in lesion locations of control subjects

Supplemental Figure 5: Sensitivity and specificity of the SE network compared to maps of brain structure, metabolism, neurotransmitter receptors, and gene expression

Supplementary Figure 6: Prediction of SE vs. controls in validation cohort, subgroups with cortical or subcortical lesion involvement

Supplementary Figure 7: Prediction of SE vs. control in the validation cohort, excluding lesions within the pulvinar or precuneus

Supplementary Figure 8: Prediction of SE vs. controls in the validation cohort, using SE networks derived from cases with different locations of seizure onset

Supplementary Figure 9: Prediction of SE vs. control in the validation cohort, using SE networks derived from cases with different SE semiology

Supplemental Figure 10: Prediction of SE vs controls in the validation cohort, using sensitivity maps and average brain connectivity (Sum of C)

Supplemental Figure 11: Prediction of SE vs controls in the validation cohort, using specificity maps controlling for average brain connectivity (Sum of C)

**Supplementary Tables:**

Supplementary Table 1: Patient demographics and lesion etiologies – Discovery cohort

Supplementary Table 2: Normative brain maps

Supplementary Table 3: Lesion overlap with cortical lobes, vascular territories, and subcortical regions – Discovery cohort

Supplementary Table 4: Prediction of SE vs. control in the validation cohort, using logistic regression models including lesion overlap with cortical lobes and vascular territories

Supplementary Table 5: Patient demographics and lesion etiologies - Validation cohort

### Supplementary Results:

#### Supplementary Results A: The SE network is robust across different control and subgroup analyses and is independent of properties of the human connectome

To test the robustness of the SE network in predicting SE-related lesion locations, we performed several control and subgroup analyses. We found that PMA overlap with the SE network was an independent predictor of SE, controlling for intersection with cortical lobes as well as vascular territories (**Supplementary Table 4**). Results were not driven by cortical or subcortical involvement as prediction of SE in the validation cohort remained robust in subgroups of patients with PMA involving either the cortex, subcortex or in patients with both cortical and subcortical involvement (**Supplementary Fig. 5**). Results were also not exclusive to lesions in the pulvinar or precuneus as predictive capacity was retained after excluding lesions involving these regions, suggesting connectivity of the lesion is key (**Supplementary Fig. 6**). The SE network was also not exclusive to any focal seizure onset zone as regenerating the network using subgroups of cases with either frontal, temporal, or posterior cortex seizure onset (as determined by scalp EEG) resulted in similar SE network topographies; each of which was able to predict SE in the validation cohort with similar accuracy (**Supplementary Fig. 7**). These findings suggest involvement of a unifying seizure network in SE independent of focal seizure onset zone. A similar result was found for convulsive versus non-convulsive SE semiologies (**Supplementary Fig. 8**). While lesion network maps may share spatial properties due to repeated sampling of average brain connectivity in the human connectome,(*1*) we demonstrate that the topography of (and peaks within) the network are specific to SE and independent of properties of the human connectome (i.e. Sum of C(*1*) **Supplementary Fig. 9 and 10**). More importantly, lesion overlap with the SE network was specific to and more predictive of SE in an independent validation cohort (AUC 90.6%) compared to multiple measures of average brain connectivity (AUC 37-42%).

### Supplemental Figures:

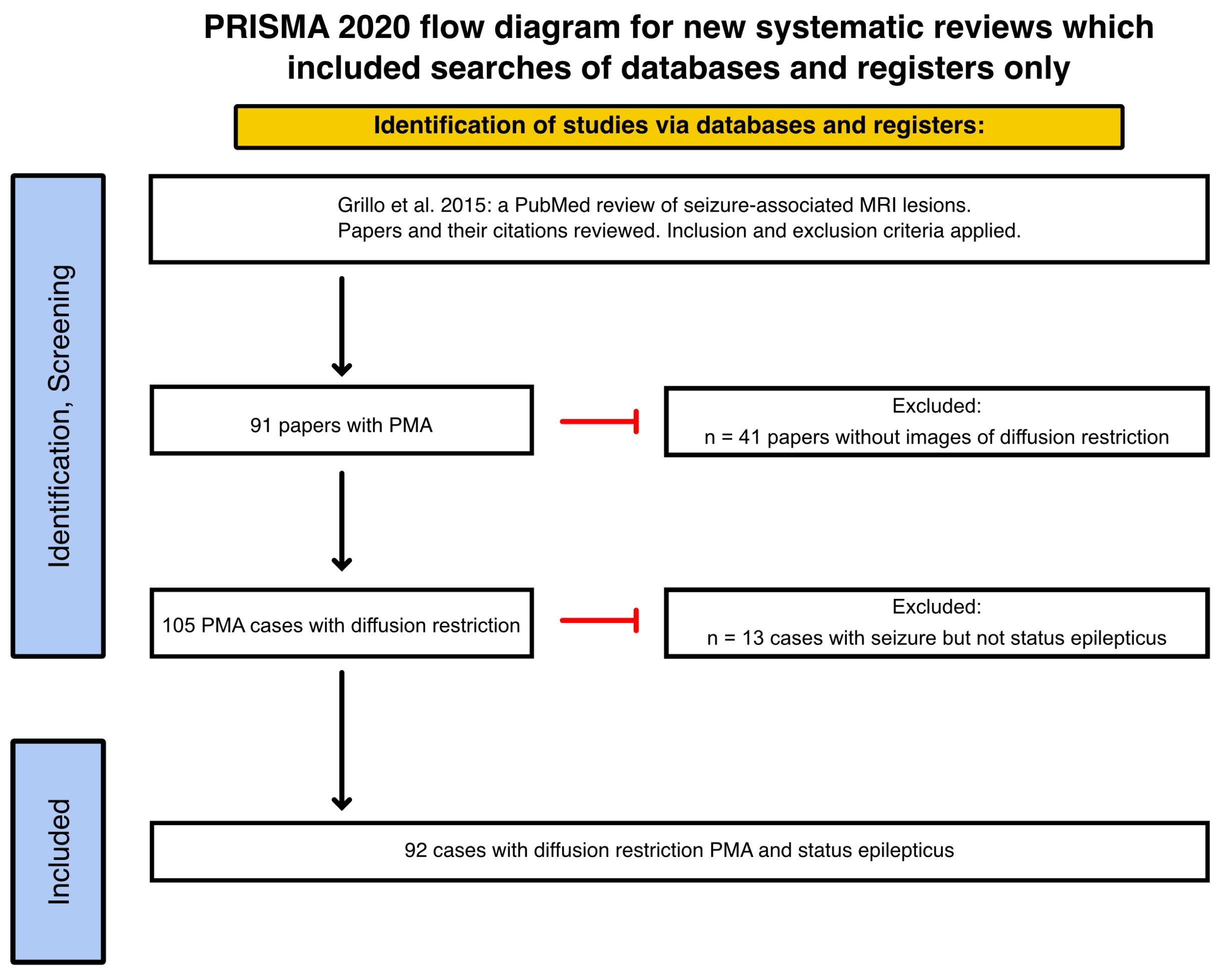


**Supplementary Figure 1: PRISMA search strategy**. From 91 papers featuring cases of peri-ictal MRI abnormalities (PMA) collated by a previously published systematic review,(*2*) 105 cases had peri-ictal diffusion restriction on MRI. From these 105 cases, 92 were diagnosed with status epilepticus per 2015 ILAE guidelines(*3*) and/or explicitly mentioned a diagnosis of 'status epilepticus' in the case report.

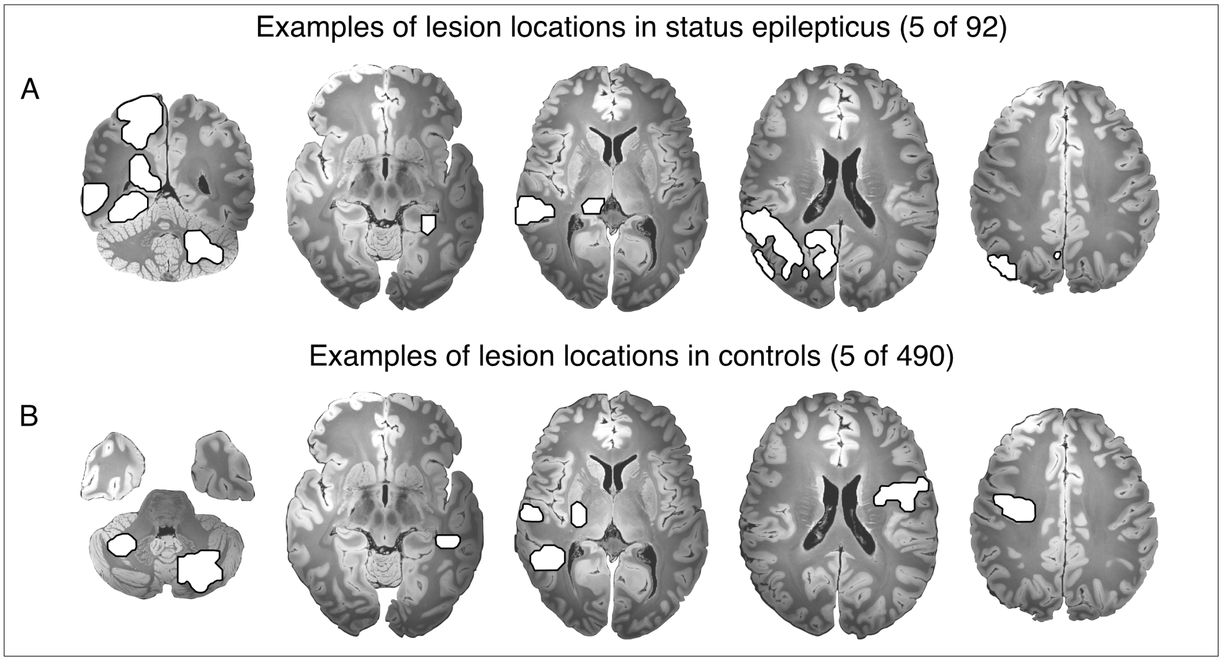


**Supplementary Figure 2. Status epilepticus and control brain lesions**. Examples of the neuroanatomical distribution of diffusion restricted brain lesions (white outlines) in patients with status epilepticus (SE) (**A**), also termed peri-ictal MRI abnormalities (PMA), as well as examples of the neuroanatomical distribution of diffusion restricted brain lesions in a control cohort of patients with stroke (**B**).

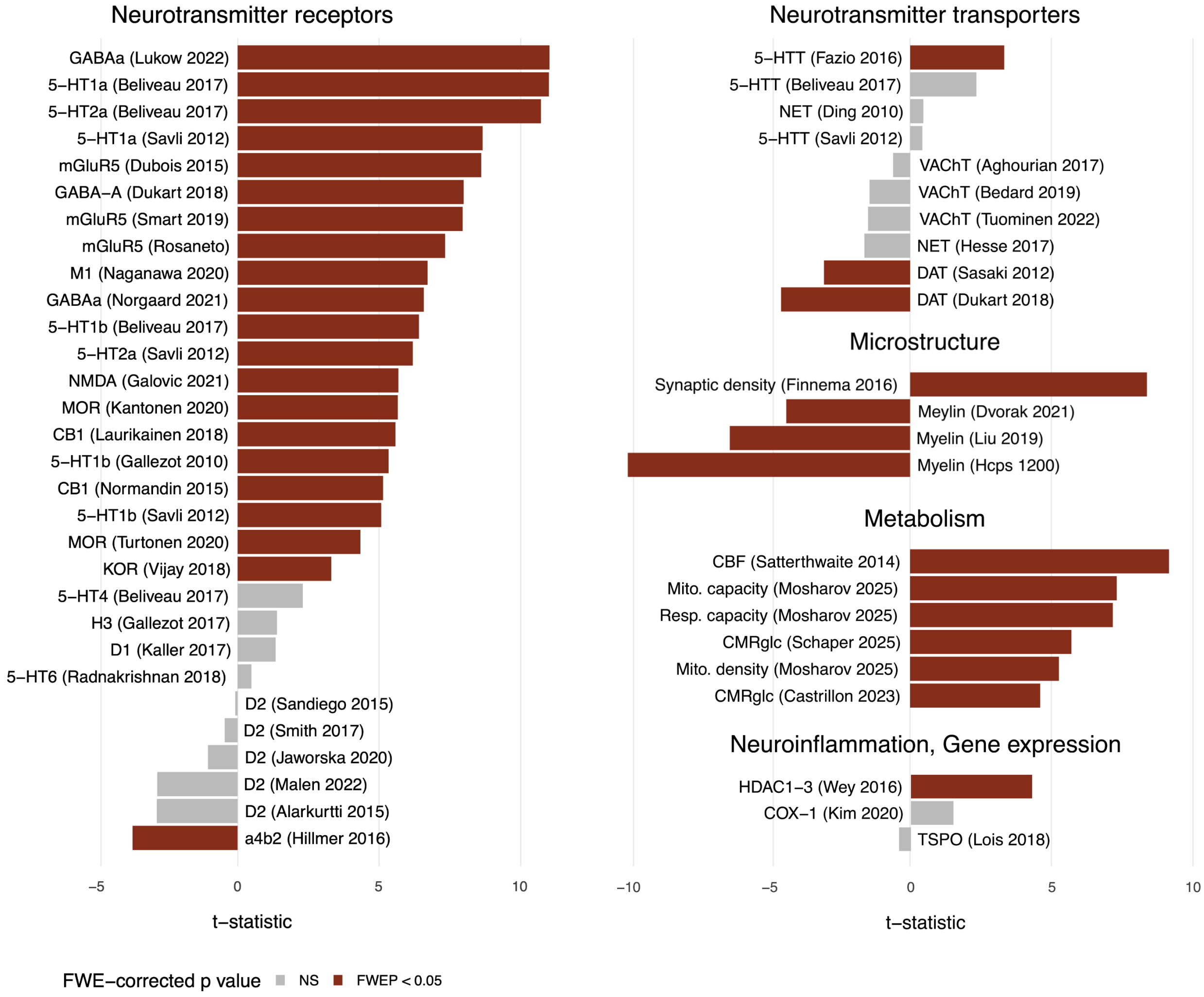


**Supplemental Figure 3: Lesion overlap of SE vs. controls in the discovery cohort on maps of brain structure, metabolism, and neurotransmitter receptors.** Lesion overlap of SE vs. control lesion locations on each of the 52 normative brain maps. From each category (neurotransmitter receptors or transporters, microstructure, metabolism, and neuroinflammation) we highlighted the most relevant non-overlapping maps with the largest effect size, resulting in the top 10 candidate maps presented in the main article. Here, we present the full results, t- and p-value for completeness. T- and p-value are presented for each individual reference map after familywise error correction for multiple comparisons across 52 maps.

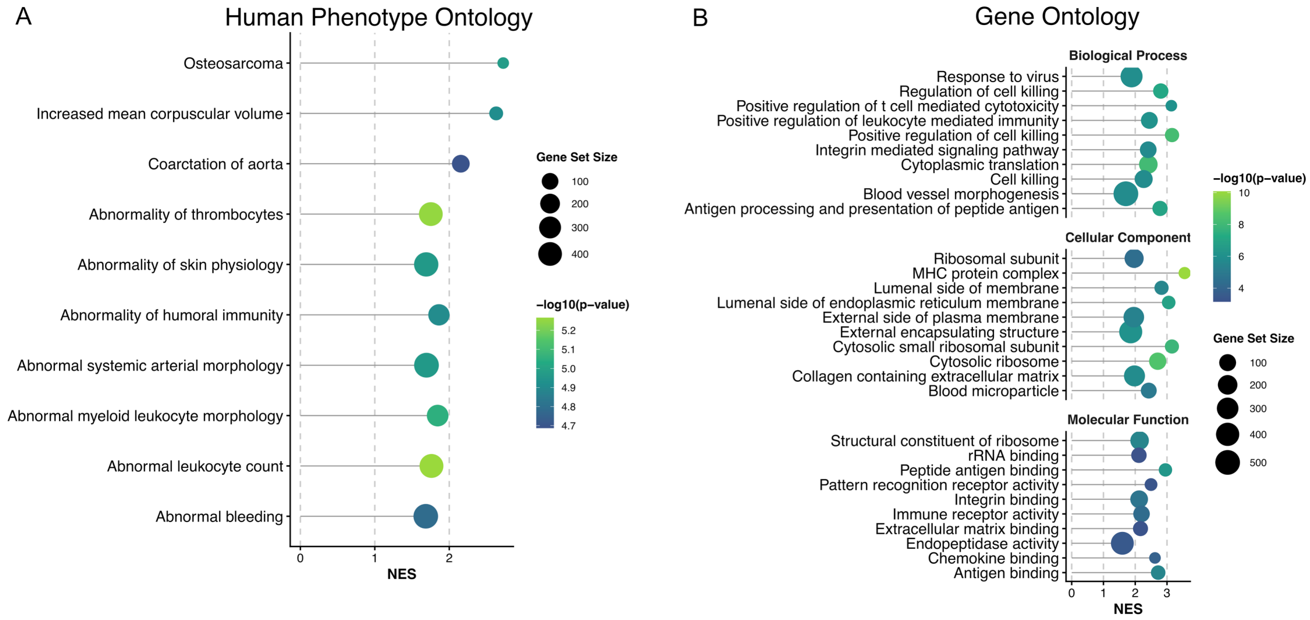


**Supplemental Figure 4: Gene expression patterns enriched in lesion locations of control subjects.** We pooled the discovery and validation cohorts and compared lesion overlap of SE and control lesions on each individual gene map (n = 15,633, thresholded z > 0). All genes 15,633 were then ranked based on effect size (highest t-value) in distinguishing between groups. To identify the gene expression patterns enriched in lesion locations of control subjects, we performed GSEA with adjusted settings (gene set size = 20-500) on the inverse of this rank list of t-values using the Human Phenotype Ontology (HPO) and Gene Ontology (GO) databases from GSEA. In contrast to SE lesion locations, control lesion locations (including strokes, tumors, and encephalitis) showed enrichment in non-neuronal processes such as development and remodeling of the vasculature, immune and extracellular matrix components, vascular and inflammatory pathologies, as well as pathways related to cytotoxicity and ribosomal proteins.

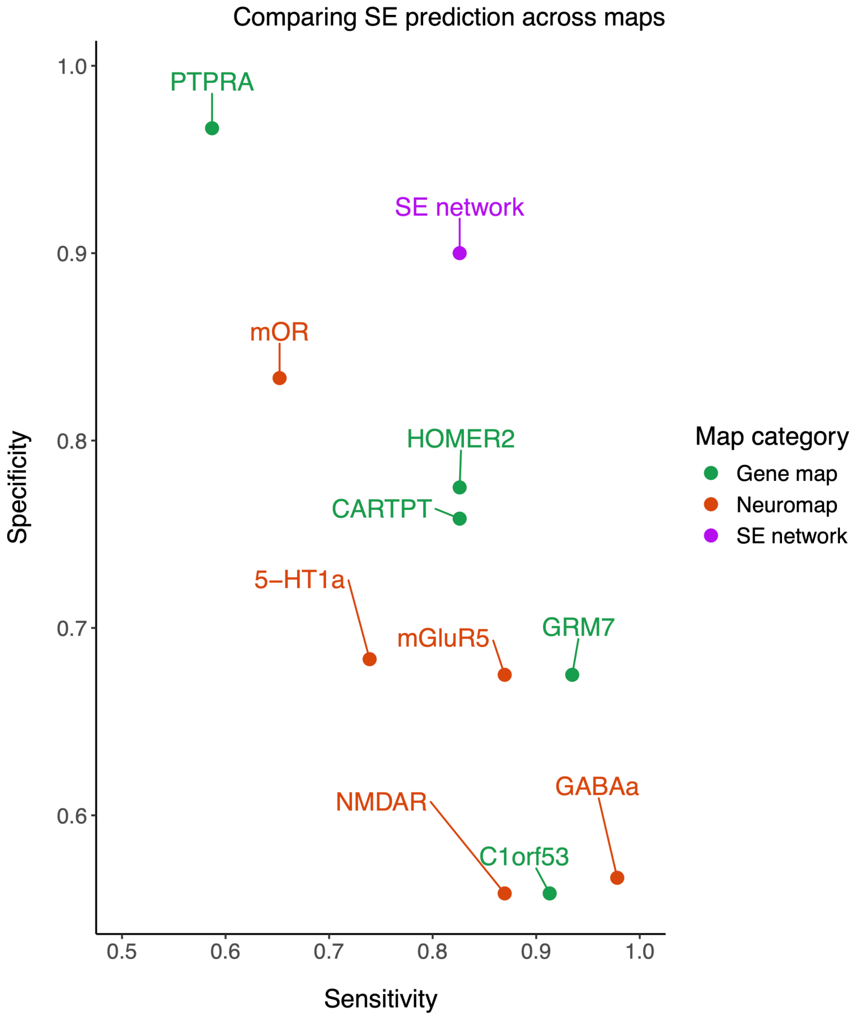


**Supplementary Figure 5: Sensitivity and specificity of the SE network compared to maps of brain structure, metabolism, neurotransmitter receptors, and gene expression.** To compare predictive value across reference maps, we computed the Youden index defined as the point on the ROC curve which maximizes the sum of sensitivity and specificity for each map. The SE network had a maximum balanced sensitivity of 82.6% and specificity of 90.0%, while other maps had notably lower specificity.

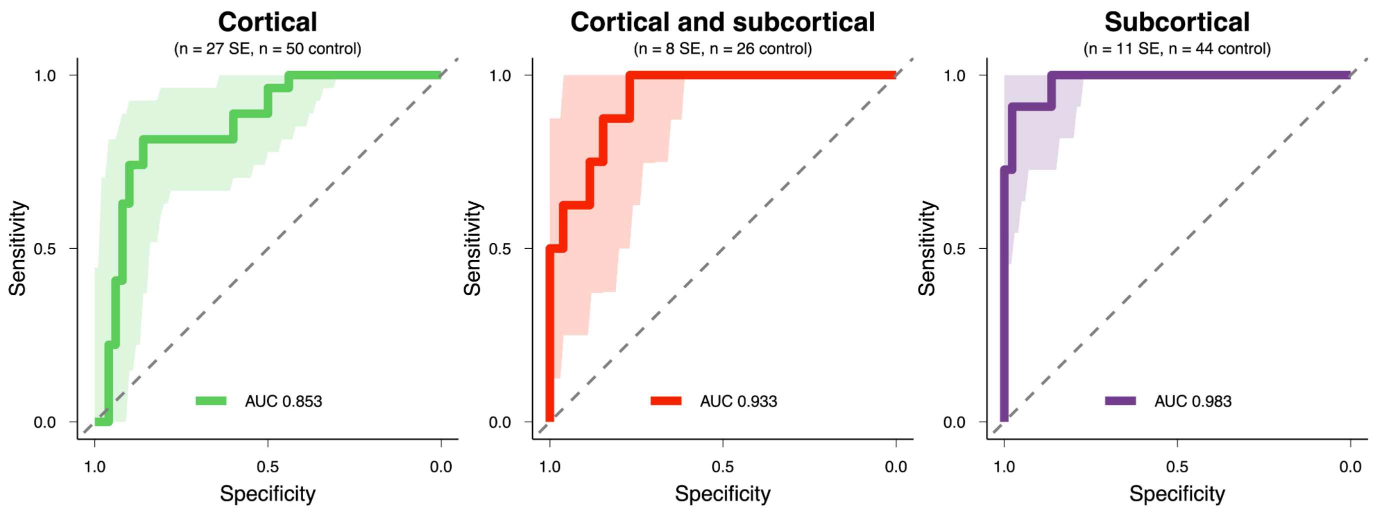


**Supplementary Figure 6: Prediction of SE vs. controls in validation cohort, subgroups with cortical or subcortical lesion involvement.** Prediction of SE vs control lesions in the validation cohort was independent of whether the lesion involved exclusively cortical or subcortical, or both cortical and subcortical brain regions. ROC were bootstrapped with 1,000 iterations to generate the 95% confidence intervals (shaded regions).

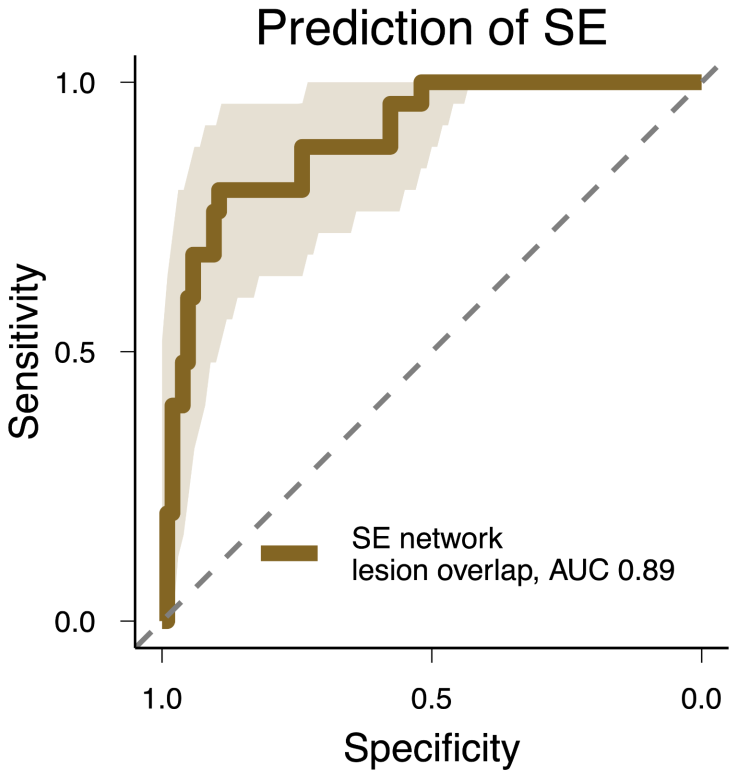


**Supplementary Figure 7: Prediction of SE vs. control in the validation cohort, excluding lesions within the pulvinar or precuneus.** Lesion overlap with the SE network remained a highly significant ($p < 0.001$) predictor of SE after removal of lesions intersecting either the pulvinar or precuneus from the validation cohort. Prediction of SE within this subgroup of patients was equally strong to prediction of the full validation cohort (DeLong test; D = 0.27167, df = 252.25, $p$ = 0.7861). ROC were bootstrapped with 1,000 iterations to generate the 95% confidence intervals (shaded regions).

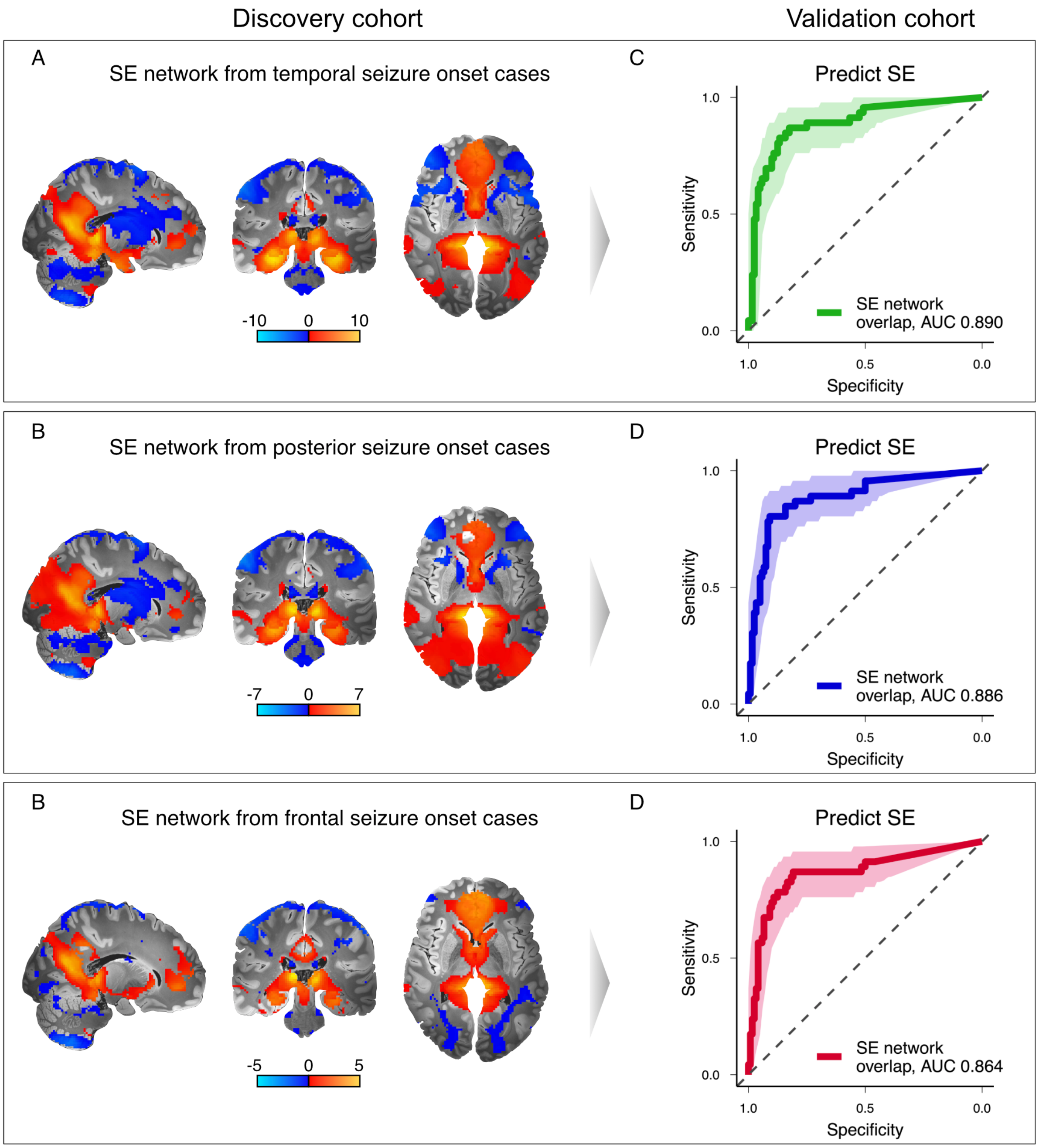


**Supplementary Figure 8: Prediction of SE vs. controls in the validation cohort, using SE networks derived from cases with different locations of seizure onset.** To test whether the SE network was independent of focal seizure onset location, we regenerated the SE network in subgroups of patients with frontal, temporal, or posterior (parietal and/or occipital) lobe seizure onset (as defined by scalp EEG). Lesion overlap with each of these variations of the SE network remained highly predictive of SE in the validation cohort, suggesting the SE network is not exclusive to any seizure onset location. Pairwise comparisons (DeLong's test) of the predictive capacity of these variations of the SE network showed no significant different across maps. ROC were bootstrapped with 1,000 iterations to generate the 95% confidence intervals (shaded regions).

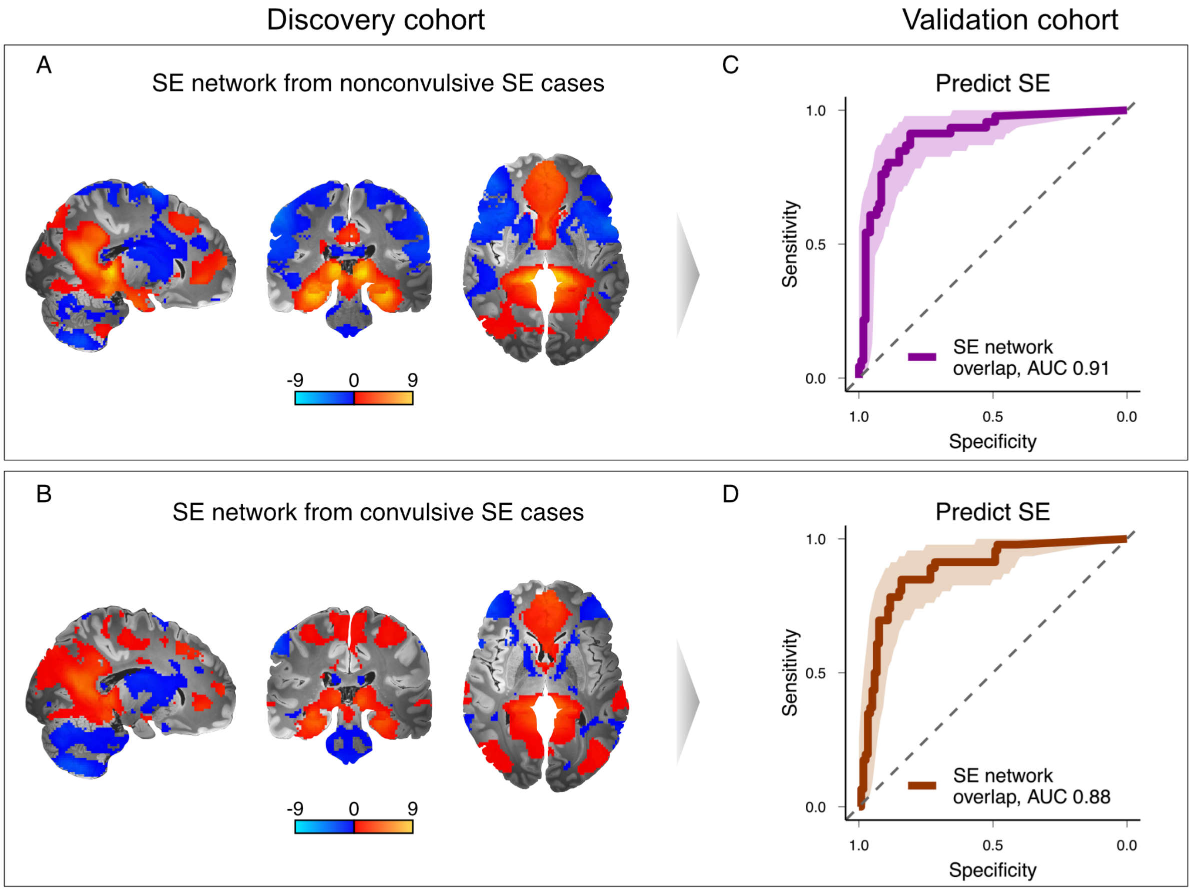


**Supplementary Figure 9: Prediction of SE vs. control in the validation cohort, using SE networks derived from cases with different SE semiology.** To test whether the SE network was independent of SE semiology, we regenerated the SE network in subgroups of patients with either convulsive or nonconvulsive SE. Lesion overlap with these variations of the SE network remained highly predictive of SE in the validation cohort. Again, there were no significant differences in the predictive capacity of either SE network variation by DeLong's test. ROC were bootstrapped with 1,000 iterations to generate the 95% confidence intervals (shaded regions).

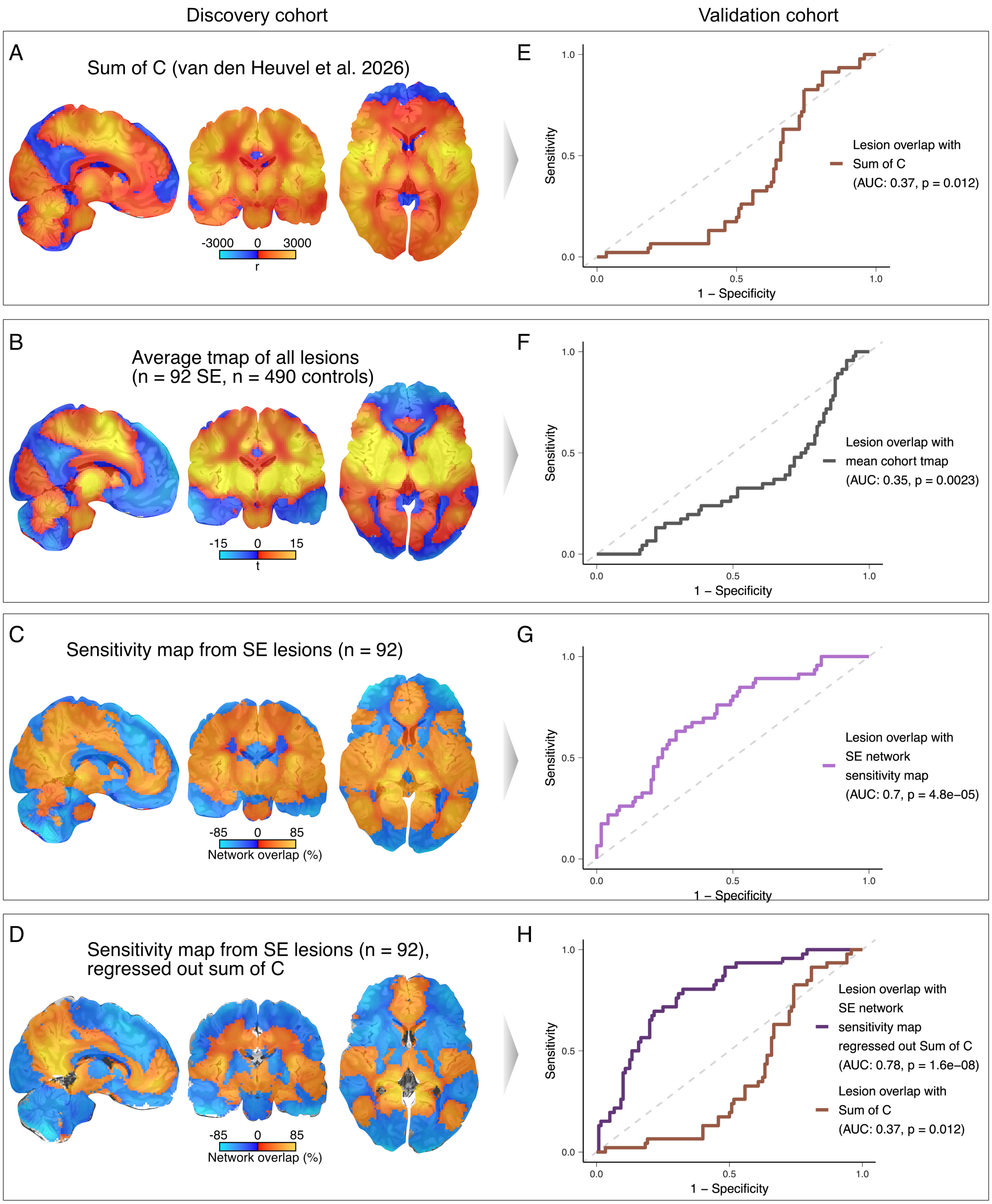


**Supplemental Figure 10: Prediction of SE vs controls in the validation cohort, using sensitivity maps and average brain connectivity (Sum of C).** van den Heuvel et al.(*1*) recently proposed that lesion network maps may share spatial properties due average brain connectivity in the human connectome (Sum of C, **A**) Indeed, when averaging the individual subject's lesion network maps of all SE and control subjects together into a single map, a connectivity profile similar to Sum of C (R = 0.69, **B**) is generated. This is consistent with the fact that lesion network mapping samples properties of the human connectome. However, it is important to emphasize that lesion network mapping does not randomly sample the human connectome but rather tests whether there is a sensitive and specific connectivity profile that is common across lesions associated with a clinical syndrome versus lesions not associated with that syndrome. In SE, sensitivity analyses show that SE lesions sample a consistent common connectivity profile that differs from Sum of C (**C**) and remains unchanged after regressing out Sum of C from each individual subject's lesion network map (**D**). More importantly, lesion overlap with any variation of the sensitivity map was a better predictor of SE in the independent validation cohort (AUC = 0.70 – 0.78, **E-H**) compared to the Sum of C map (AUC = 0.37, **E,H**). These findings suggest sensitivity analyses in lesion network mapping provide unique and relevant information not captured by average brain connectivity.

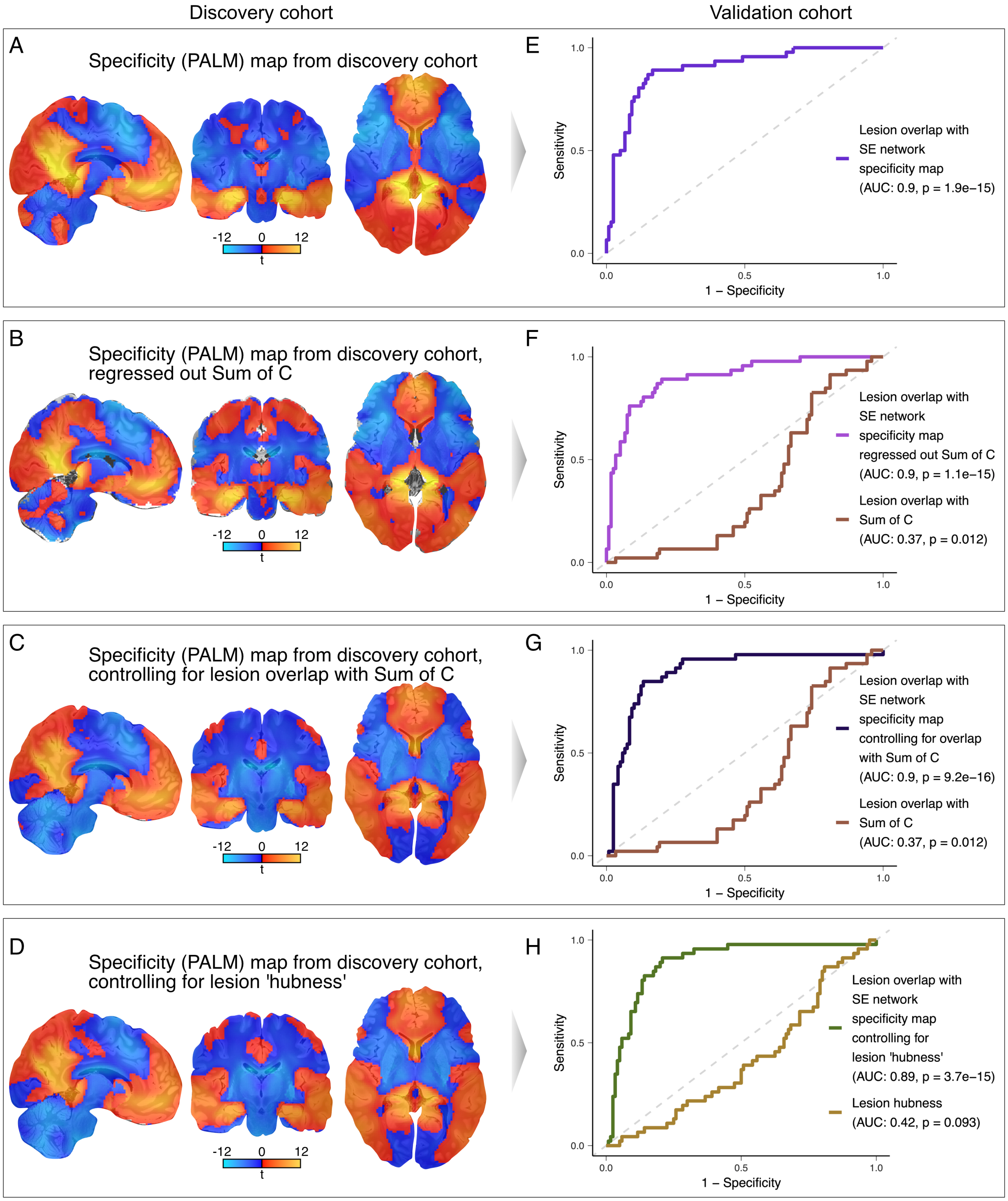


**Supplementary Figure 11: Prediction of SE vs controls in the validation cohort, using specificity maps controlling for average brain connectivity (Sum of C).** Specificity analyses were performed using a voxel-wise permutation-based t-test, which demonstrated that SE lesions sample a specific connectivity profile that differs from control lesions (**A**). This specificity map of SE versus control lesions (**A**) differs from the Sum of C map and remains unchanged after either regressing out Sum of C (**B**), controlling for lesion overlap with Sum of C (**C**) or controlling for lesion hubness (**D**) as a covariate in the voxel-wise permutation-based t-test. Lesion hubness was computed for each individual lesion as a measure of average brain connectivity (centrality) by summing the R values outside a given lesion mask and divide by lesion volume, consistent with how the Sum of C map was calculated in van den Heuvel et al. 2026.(*1*) More importantly, lesion overlap with any variation of the specificity map of lesion network mapping was a better predictor of SE in the independent validation cohort (AUC = 0.90), **E-H**) compared to any measure of average brain connectivity (Sum of C, AUC = 0.37 – 0.42, **F-H**). These findings suggest specificity analyses in lesion network mapping provide unique and relevant information not captured by average brain connectivity.

### Supplemental Tables:

**Supplementary Table 1: Patient demographics and lesion etiology – discovery cohort**

| **Characteristic** | **SE**<br>N = 92[1] | **Control**<br>N = 490[1] |
|---|---|---|
| Age | 52 (35, 71) | 65 (50, 80) |
| Sex | | |
| F | 39 (42%) | 303 (62%) |
| M | 42 (46%) | 187 (38%) |
| Unreported | 11 (12%) | 0 (0%) |
| Semiology Convulsive | | |
| Yes | 45 (49%) | N/A |
| No | 39 (42%) | N/A |
| Unreported | 8 (8.7%) | N/A |
| N/A | 0 (0%) | 490 (100%) |
| EEG Onset Focal | | |
| Focal | 62 (67%) | N/A |
| Unreported | 19 (21%) | N/A |
| Generalized | 11 (12%) | N/A |
| N/A | 0 (0%) | 490 (100%) |
| Etiology | | |
| Cerebrovascular | 22 (24%) | 490 (100%) |
| Toxic-Metabolic | 21 (23%) | 0 (0%) |
| Unknown/Unreported | 20 (22%) | 0 (0%) |

**Supplementary Table 1: Patient demographics and lesion etiology – discovery cohort**

| **Characteristic** | **SE**<br>N = 92[1] | **Control**<br>N = 490[1] |
|---|---|---|
| Chronic Epilepsy of Unknown Etiology | 13 (14%) | 0 (0%) |
| Neoplasm | 10 (11%) | 0 (0%) |
| Structural/Neurodevelopmental disorder | 4 (4.3%) | 0 (0%) |
| Other | 2 (2.2%) | 0 (0%) |

[1]Median (Q1, Q3); n (%)

**Supplementary Table 2: Patient demographics and lesion etiology - validation cohort**

| Characteristic | SE<br>N = 46[1] | Control<br>N = 120[1] |
|---|---|---|
| Sex | | |
| M | 20 (43%) | 66 (55%) |
| F | 26 (57%) | 54 (45%) |
| Age | 68 (55, 78) | 70 (58, 77) |
| Etiology | | |
| Cerebrovascular disease | 15 (33%) | 108 (90%) |
| Intracranial tumor | 7 (15%) | 8 (6.7%) |
| Head trauma | 7 (15%) | 0 (0%) |
| Cryptogenic/NR | 5 (11%) | 0 (0%) |
| Infectious/Inflammatory | 5 (11%) | 0 (0%) |
| Metabolic/Toxidrome | 4 (8.7%) | 0 (0%) |
| Hypoxic encephalopathy | 0 (0%) | 2 (1.7%) |
| Kikuchi-Fujimoto disease | 1 (2.2%) | 0 (0%) |
| Multiple sclerosis | 0 (0%) | 1 (0.8%) |
| Neurodegenerative | 1 (2.2%) | 0 (0%) |
| Transient global amnesia | 0 (0%) | 1 (0.8%) |
| Withdrawal of ASM | 1 (2.2%) | 0 (0%) |

[1]n (%); Median (Q1, Q3)

**Supplementary Table 3: Lesion overlap with cortical lobes, vascular territories, and subcortical regions – Discovery cohort.**

| | Brain region | SE N = 92[1] | Control N = 490[1] | Chi-square | FWE-corrected p-value |
|---|---|---|---|---|---|
| Laterality | Right hemisphere | 64 (70%) | 282 (58%) | 4.64 | 0.998 |
| | Left hemisphere | 64 (70%) | 288 (59%) | 3.77 | >0.999 |
| Cortical, subcortical | Cortical | 75 (82%) | 289 (59%) | 16.80 | 0.002 |
| | Subcortical | 92 (100%) | 481 (98%) | 1.72 | >0.999 |
| Lobe | Occipital lobe | 47 (51%) | 87 (18%) | 48.56 | <0.001 |
| | Temporal lobe | 46 (50%) | 99 (20%) | 36.76 | <0.001 |
| | Parietal lobe | 53 (58%) | 181 (37%) | 13.77 | 0.008 |
| | Frontal lobe | 41 (45%) | 183 (37%) | 1.70 | >0.999 |
| | Insular lobe | 23 (25%) | 96 (20%) | 1.39 | >0.999 |
| Vascular territory | PCA | 76 (83%) | 217 (44%) | 45.51 | <0.001 |
| | ACA | 39 (42%) | 167 (34%) | 2.34 | >0.999 |
| | MCA | 74 (80%) | 385 (79%) | 0.16 | >0.999 |
| | Infratentorial vasculature | 20 (22%) | 98 (20%) | 0.14 | >0.999 |
| Region | Limbic system | 58 (63%) | 67 (14%) | 111.95 | <0.001 |
| | Pulvinar | 32 (35%) | 39 (8.0%) | 52.03 | <0.001 |
| | Precuneus | 25 (27%) | 24 (4.9%) | 49.85 | <0.001 |
| | Hippocampus | 21 (23%) | 17 (3.5%) | 47.55 | <0.001 |
| | Amygdala | 8 (8.7%) | 6 (1.2%) | 18.42 | 0.002 |
| | Thalamus | 34 (37%) | 98 (20%) | 12.70 | 0.015 |
| | Corpus callosum | 20 (22%) | 54 (11%) | 8.02 | 0.187 |
| | Basal ganglia | 15 (16%) | 142 (29%) | 6.32 | 0.409 |
| | Cerebellum | 11 (12%) | 66 (13%) | 0.15 | >0.999 |

[1]n (%)

[2]Pearson's Chi-squared test

[3]Bonferroni correction

| Supplementary Table 4: Normative brain maps. | | |
|---|---|---|
| | **Biomarker** | **Published article DOI** |
| **Neurotransmitter Transporter** | | |
| Acetylcholine transporter | VAChT | 10.1038/mp.2017.183 |
| | VAChT | 10.1016/j.sleep.2018.12.020 |
| | VAChT | 10.1038/s41593-022-01186-3 |
| Dopamine transporter | DAT | 10.2967/jnumed.111.101626 |
| | DAT | 10.1038/s41598-018-22444-0 |
| Norepinephrine transporter | NET | 10.1002/syn.20696 |
| | NET | 10.1007/s00259-016-3590-3 |
| Serotonin transporter | 5-HTT | 10.1523/JNEUROSCI.2830-16.2016 |
| | 5-HTT | 10.1016/j.neuroimage.2016.03.019 |
| | 5-HTT | 10.1016/j.neuroimage.2012.07.001 |
| **Neurotransmitter Receptor** | | |
| Acetylcholine receptor | M1 mAChR | 10.2967/jnumed.120.246967 |
| | Aβ42 | 10.1016/j.neuroimage.2016.07.026 |
| Cannabinoid receptor | CB1 | 10.1371/journal.pone.0060231 |
| | CB1 | 10.1038/jcbfm.2015.46 |
| Dopamine receptor | D1 | 10.1007/s00259-017-3645-0 |
| | D2 | 10.1038/jcbfm.2015.53 |
| | D2 | 10.1038/s41386-020-0662-7 |
| | D2 | 10.1016/j.neuroimage.2022.119149 |
| | D2 | 10.1038/jcbfm.2014.237 |
| | D2 | 10.1177/0271678X17737693 |
| GABA receptor | GABAA | 10.1016/j.neuroimage.2021.117878 |
| | GABAA | 10.1038/s41598-018-22444-0 |
| | GABAA α-5 subunit | 10.1038/s42003-022-03268-1 |
| Glutamate receptor | mGluR5 | 10.1007/s00259-015-3167-6 |
| | mGluR5 | 10.1038/s41593-022-01186-3 |
| | mGluR5 | 10.1007/s00259-018-4252-4 |
| | NMDAR | 10.1016/j.neuroimage.2021.118194 |

| **Supplementary Table 4: Normative brain maps.** | | |
|---|---|---|
| | **Biomarker** | **Published article DOI** |
| Histamine receptor | H3 | 10.1038/jcbfm.2009.195 |
| Serotonin receptor | 5-HT1a | 10.1523/JNEUROSCI.2830-16.2016 |
| | 5-HT1a | 10.1016/j.neuroimage.2012.07.001 |
| | 5-HT1b | 10.1523/JNEUROSCI.2830-16.2016 |
| | 5-HT1b | 10.1038/jcbfm.2009.195 |
| | 5-HT1b | 10.1016/j.neuroimage.2012.07.001 |
| | 5-HT2a | 10.1523/JNEUROSCI.2830-16.2016 |
| | 5-HT2a | 10.1016/j.neuroimage.2012.07.001 |
| | 5-HT4 | 10.1523/JNEUROSCI.2830-16.2016 |
| | 5-HT6 | 10.2967/jnumed.117.206516 |
| κ-opioid receptor | KOR | 10.1038/s41386-018-0199-1 |
| μ-opioid receptor | MOR | 10.1016/j.neuroimage.2020.116922 |
| | MOR | 10.1016/j.bpsc.2020.10.013 |
| **Neuroinflammation** | | |
| Neuroinflammatory response | COX-1 | 10.1007/s00259-020-04855-2 |
| | TSPO | 10.1021/acschemneuro.8b00072 |
| **Microstruture** | | |
| Myelin | Myelin water fraction | 10.1038/s41598-020-79540-3 |
| **Microstructure** | | |
| Myelin | Myelin water fraction | 10.1111/jon.12657 |
| | T1w/T2w ratio | https://www.humanconnectome.org/study/hcp-young-adult/article/s1200-group-average-data-release |
| Synaptic neurotransmission | SV2A | 10.1177/0271678X17724947 |
| **Metabolism** | | |
| Metabolic markers | Cerebral blood flow | 10.1073/pnas.1400178111 |
| | Glucose metabolism | 10.1126/sciadv.adi7632 |
| | Mitochondrial Density | 10.1038/s41586-025-08740-6 |

| Supplementary Table 4: Normative brain maps. | | |
|---|---|---|
| | **Biomarker** | **Published article DOI** |
| | Oxidative phosphorylation enzyme activity | 10.1038/s41586-025-08740-6 |
| | Tissue respiratory capacity / mitochondrial density | 10.1038/s41586-025-08740-6 |
| | **Epigenetic marker** | |
| Epigenetic markers | HDAC1, 2, 3 | 10.1126/scitranslmed.aaf7551 |

**Supplementary Table 5: Prediction of SE vs. controls in the validation cohort, using logistic regression models including lesion overlap with cortical lobes and vascular territories.**

| | SE Network | PCA territory | Frontal lobe | Temporal lobe | Parietal lobe | Occipital lobe | All lobes | SE + PCA | SE + Lobes |
|---|---|---|---|---|---|---|---|---|---|
| Overlap with SE network | 6.26 | | | | | | | 6.07 | 6.46 |
| | (3.60 - 12.03), p = <0.01 | | | | | | | (3.47 - 11.79), p = <0.01 | (3.58 - 13.10), p = <0.01 |
| Voxels in PCA territory | | 1.33 | | | | | | 1.08 | |
| | | (0.97 - 1.96), p = 0.09 | | | | | | (0.76 - 1.53), p = 0.64 | |
| Voxels in Frontal Lobe | | | 1.23 | | | | 1.06 | | 1.68 |
| | | | (0.89 - 1.86), p = 0.23 | | | | (0.41 - 2.30), p = 0.88 | | (0.69 - 3.86), p = 0.20 |
| Voxels in Temporal Lobe | | | | 1.47 | | | 2.78 | | 1.53 |
| | | | | (1.06 - 2.22), p = 0.04 | | | (1.41 - 6.13), p = <0.01 | | (0.74 - 3.32), p = 0.24 |
| Voxels in Parietal Lobe | | | | | 1.35 | | 12.49 | | 16.87 |
| | | | | | (0.97 - 2.21), p = 0.12 | | (2.32 - 105.30), p = <0.01 | | (2.54 - 225.46), p = <0.01 |
| Voxels in Occipital Lobe | | | | | | 1.15 | 0.05 | | 0.04 |
| | | | | | | (0.82 - 1.61), p = 0.38 | (0.00 - 0.37), p = <0.01 | | (0.00 - 0.35), p = <0.01 |
| AIC | 134.3 | 196.8 | 198.3 | 194.7 | 196.8 | 199.2 | 188.4 | 136.1 | 129.0 |
| RMSE | 0.34 | 0.44 | 0.45 | 0.44 | 0.44 | 0.45 | 0.42 | 0.34 | 0.32 |
| McFadden pseudo-$R^2$ | 0.335 | 0.016 | 0.008 | 0.027 | 0.016 | 0.004 | 0.090 | 0.336 | 0.403 |

Note: Results display Odds Ratios (OR) and their 95% Confidence Intervals (CI) for scaled lesion network overlap and lesion voxel overlap. ORs are calculated by exponentiating the logistic regression coefficients.

**Supplementary Table 5:** Logistic regression was performed to predict SE in the validation cohort using the following (scaled) independent variables: lesion overlap with the SE network, cortical lobes, or posterior cerebral artery (PCA) vascular territory. Lesion overlap with the SE network was an independent predictor of SE in a model including PCA overlap or any cortical lobe. Lesion overlap with the SE network was also the strongest single predictor of SE, while controlling for any other variable.
